# Resolving human α *versus* β cell fate allocation for the generation of stem cell-derived islets

**DOI:** 10.1101/2024.06.20.599862

**Authors:** Melis Akgün Canan, Corinna Cozzitorto, Michael Sterr, Lama Saber, Eunike S.A. Setyono, Xianming Wang, Juliane Merl-Pham, Tobias Greisle, Ingo Burtscher, Heiko Lickert

## Abstract

Generating stem cell-derived glucagon-producing α (SC-α cells) and insulin-producing β cells (SC-β cells) allows to engineer an *in vitro* biomimetic of the islet of Langerhans, the micro-organ controlling blood glucose, however, there is still a major knowledge gap in the mode and mechanism by which human SC-α and β cells are specified. Mouse studies postulated that Aristaless Related homeobox (Arx) and Paired box 4 (Pax4) transcription factors cross-inhibit each other in endocrine progenitors to promote α or β cell fate allocation, respectively. To test this model in human, we generated an *ARX^CFP/CFP^; PAX4^mCherry/mCherry^* double knock-in reporter induced pluripotent stem cell (iPSC) line to combine time-resolved cell lineage labeling with high-resolution single cell multiomic analysis. Strikingly, lineage labelling and tracing, proteomic and gene regulatory network (GRN) analysis and potency assays revealed a human specific mode and regulatory logic of α *versus* β cell fate allocation. Importantly, pharmacological perturbation using drugs previously proposed to trigger α-to-β cell transdifferentiation or identified via our GRN analysis led to enhanced endocrine induction and directed α vs β cell fate commitment. Thus, shedding light on basic mechanisms of endocrine induction and fate segregation not only paves the way to engineer islets from pluripotent stem cells, but also has broader implications for cell-replacement therapy, disease modelling and drug screening.

## INTRODUCTION

Diabetes is a metabolic disease characterized by impaired blood glucose homeostasis due to insufficient insulin levels. Causes are immune-dependent destruction of insulin-producing β cells (type 1 diabetes – T1D) or β cell dysfunction (type 2 diabetes – T2D)^1,2^. Depending on the species, β cells make up between 60% (human and non-human primates) to 80% (mouse) of the hormone-producing endocrine compartment of the pancreas, the islet of Langerhans. Other cellular components of pancreatic islets are glucagon-producing α cells, somatostatin-producing δ cells, together with ε and PP cells producing ghrelin and pancreatic polypeptide, respectively^3–5^.

Islet transplantation is an effective treatment to reach independence from exogenous insulin, but availability of cadaveric islets from human donors in scarce^6^. Within the search for possible replacements for lost or dysfunctional β cells, efforts over the years focused on the establishment of differentiation protocols that could provide an unlimited source of stem cell-derived β cells (SC-β cells) for cell therapy^7,8^. Stepwise differentiation protocols mimic pancreatic development, progressing through anterior definitive endoderm (ADE), primitive gut tube (PGT), pancreatic progenitor (PP), and endocrine progenitor (EP) stages to guide the differentiation of hormone-producing cells *in vitro* starting from diverse human pluripotent stem cell (PSC) sources^9–18^. These protocols not only heavily rely on the knowledge acquired from studying mouse pancreas development, but also use empirically tested chemicals to improve certain stages of differentiation, resulting into cell products (SC-islets) that do not completely resemble neither human nor mouse islets. In addition to generate heterogenous SC-islets comprising undesired cell types, state of the art protocols progressively lose differentiation efficiency starting from the induction of EPs to the differentiation of SC-β cells^8,19^.

To improve the differentiation efficiency towards either SC-α or β cells, a better understanding of the mechanisms of endocrine cell specification during human pancreas development is needed. In the mouse, endocrine cell fate decision towards α or β cells take place when EPs expressing the endocrine master regulator *Neurogenin 3* (*Neurog3, Ngn3*) start to express the transcription factors (TFs) *Aristaless Related homeobox* (*Arx*) or *Paired Box 4* (*Pax4*), respectively^20,21^. *Arx* knock-out (KO) mice have increased number of β cells with a near total loss of α cells, whereas KO mice for *Pax4* have increased number of α cells and virtually no β cells^20,22^. In addition, double KO mice for *Arx* and *Pax4* show virtual total loss of both α and β cells, with increased numbers of δ and PP cells^23^. These genetic studies support a model of α *versus* β cell fate decision in which Arx and Pax4 stochastically compete with each other to promote α or β cell differentiation^21,23^. However, many questions remain open, i.e. 1) How would this direct cross-regulation be achieved mechanistically? 2) What are the upstream regulators of both *ARX* and *PAX4* transcription, and consequently α vs β cell fate allocation? 3) Is this mouse-based model conserved during evolution and in human pancreas development? All these aspects have never been tested due to the lack of available tools to directly assess ARX and PAX4 activity and potency. As of the preparation of this manuscript, only three protocols have been established to direct pluripotent stem cell differentiation toward the α cell fate^24–26^, and no human model has been established to follow spatio-temporally ARX and PAX4 activity to decipher mechanistically how α vs β cell fate allocation occurs.

Studies in mouse models proposed α-to-β cell fate conversion from residual or supernumerary α cells as a potential mechanism for β cell regeneration in T1D^27–29^. In the search for potential drugs promoting this mechanism, one study showed artemether as a trigger for α-to-β cell transdifferentiation both *in vitro* and *in vivo*^30^. Since then, the effect of the antimalaria drugs Artemisinins on *Arx* and *Pax4* expression and, more in general, on pancreatic cell fate decision and regeneration is controversial. Some studies reported their impact on Arx and Pax4 expression and/or function, endocrine cell fate commitment and regeneration of β cells^30–32^, while other authors reported opposite results or no changes upon treatment^33–35^. It is important to notice that while these studies explored the effect of Artemisinins in multiple species and experimental settings, to the best of our knowledge, none of them interrogated the effect of this class of antimalaria drugs during human pancreatic endocrine development.

Here, we generated a hiPSC fluorescent double reporter line to follow ARX and PAX4 dynamics. We first used this newly established tool to study the dichotomy within the two TFs. We used multiomic and proteomic analysis together with cell potency assays to generate the first human GRN involving both *ARX* and *PAX4* and propose a new model of human α *versus* β cell fate decision. We then used the same cell line to test the effect of drugs that have been identified by our GRN analysis or being proposed as triggers of α-to-β cell transdifferentiation on α *versus* β cell fate decision. In particular, we showed that treatment with FoxO1 small molecule inhibitors increases the number of SC-β cells, and the treatment with artemether during human SC-islets differentiation promotes late PP expansion, endocrine induction and increases β cell differentiation at the expense of α cells. With the generation of a unique ARX/PAX4 hiPSC reporter line, our results shed new light on the mode of human endocrinogenesis and pave the way for drug testing to improve differentiation protocols to generate SC-islets with a defined ratio of SC-α and β cells to mimic the islet of Langerhans.

## RESULTS

### PAX4 and ARX are not co-expressed in early NEUROG3^+^ progenitors *in vitro* and *in vivo*

PAX4 and ARX TFs govern pancreatic EP lineage fate decision toward α or β cells in the mouse^21^. PAX4 promotes β cell differentiation, while ARX promotes differentiation towards the α cell lineage^20,22,23^ (Fig. 1A). This model would require *PAX4* and *ARX* to be mostly co-expressed upon induction and segregation in early EPs to allow the two proteins to act on each other *locus* and stochastically drive α or β cell fate commitment. To test if this minimal requirement is fulfilled during human pancreas development *in vivo*, we interrogated a recently published human atlas of fetal pancreas development^37^. We looked at the expression of *PAX4*, *ARX*, and *NEUROGENIN3* (*NEUROG3*), a marker of EPs and master regulator of endocrine induction, in single cell RNA-Sequencing (scRNA-Seq) datasets (Fig. 1B). This analysis revealed a transient *PAX4* co-expression with *NEUROG3* in EPs (EP1 and FEV clusters) from 8 weeks post conception (wpc), with a peak of expression in pre-β cells at 12 wpc that decreased at later stages and was not detectable in insulin-producing β cells. On the other hand, *ARX* expression was restricted to pre-α cells and pre-ε cells starting from 8 wpc, peaked around 12 wpc and was sustained at later stages in glucagon-positive α cells. This observation suggests that in human the commonly accepted mode of action of PAX4 and ARX (Fig. 1A) is likely not accurate, as the two opposing TFs are not largely co-expressed in early *NEUROG3*-positive EPs.

**Fig. 1.**
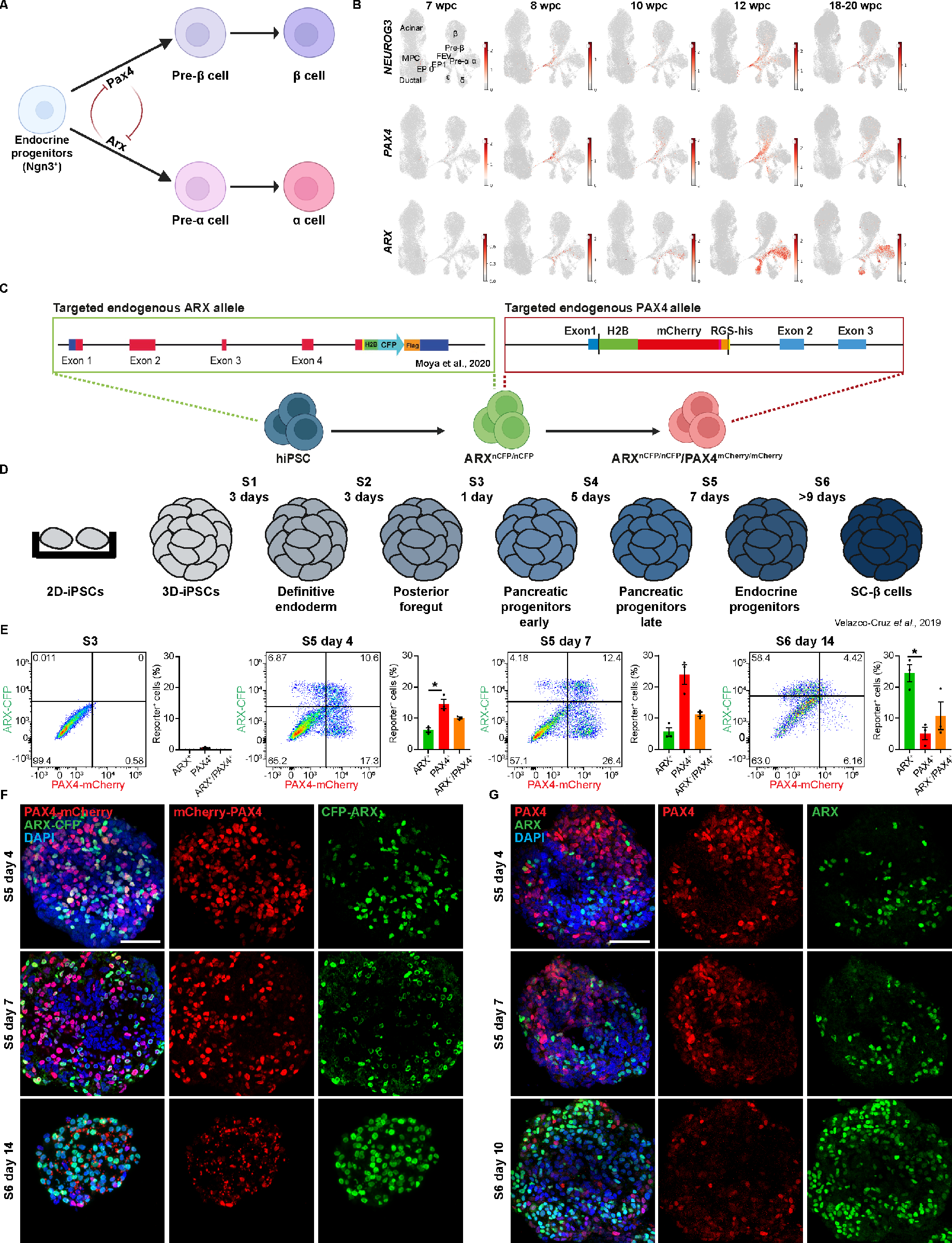
PAX4- and ARX-reporters have asynchronous expression patterns. **(A)** Postulated NGN3-PAX4-ARX signaling cascade involved in α and β cells differentiation in mouse. **(B)** Uniform Manifold Approximation and Projections (UMAPs) showing *NEUROG3, PAX4* and *ARX* mRNA expression within human pancreata at 7-, 8-, 10-, 12- and 18-20-weeks post conception (wpc). **(C)** Generation strategy of the *ARX^CFP/CFP^/PAX4^mCherry/mCherry^* reporter hiPSC line via CRISPR-Cas9-based targeting starting from the *ARX^nCFP/nCFP^* reporter cell line ^36^. H2B-mCherry was inserted in exon 1 of the endogenous *PAX4* allele together with an RGS-histidine tag. Two homozygous clones for both genetic alterations were generated, and one used for all subsequent experiments. **(D)** Differentiation protocol as in Velazco-Cruz *et al.*, 2019. **(E)** Representative flow cytometry plots and related quantifications of ARX-CPF and PAX4-mCherry reporters-positive cells during SC-β cell differentiation at PP stage (S3), mid (S5 day 4-S5d4) and end of EP stage (S5d7), and SC-β cell stage (S6d14). n=3 from distinct differentiation experiments. **(F-G)** Representative maximum intensity projections of Z-stack confocal acquisitions of immunofluorescence (IF) staining for ARX-CFP and PAX4-mCherry in ARX/PAX4 reporter clusters (F) and endogenous ARX and PAX4 in wild-type clusters (G) at the indicated stages. Data are presented as mean ± SE. One-way ANOVA with Tukey multiple comparison test. * ≤ 0.05. Scale bars 50 µm.

Due to the several limitations concerning the study of human endocrinogenesis *in vivo*, we took advantage of established *in vitro* human β cell differentiation protocols to study α *versus* β cell fate decision and to analyze the role of PAX4 and ARX in this context. We first interrogated an already published scRNA-Seq dataset of human PSC lines undergoing β cell differentiation *in vitro*^38^. We looked at the expression of *NEUROG3, PAX4*, and *ARX* in stem cell-derived EPs (stage 5 – S5). In line with our *in vivo* findings, *NEUROG3* and *PAX4* were largely co-expressed in early, mid, and late EPs, while *ARX* was expressed in α and in δ cells expressing *Somatostatin* (*SST*) and *Haematopoietically Expressed Homeobox* (*HHEX*) (Suppl. Fig. 1A-B). To temporally resolve and mechanistically understand how, and to which extent, ARX and PAX4 control α and β cell fate decision at the single cell level, we generated a homozygous human iPSC line expressing the CFP reporter for ARX and the mCherry reporter for PAX4 (*ARX^nCFP/nCFP^*; *PAX4^mCherry/mCherry^*), hereafter called “ARX/PAX4 reporter”. Using CRISPR/Cas9 engineering and our previously published *ARX^nCFP/nCFP^* reporter iPSC line^36^, we inserted the coding sequence of a nuclear tagged (histone 2B - H2B) mCherry fluorescent protein in exon 1 of the endogenous *PAX4* allele (Fig. 1C). Consequently, transcription of each TF results into polycistronic messenger RNAs (mRNAs) encoding both the TF and the fluorescent protein with a T2A sequence between them. Therefore, ARX and PAX4 proteins are not fused to the fluorescent reporters, but co-translation of TFs and reporters is equimolar. Using this strategy, we were able to simultaneous look at the rate of transcription of both genes encoding the TFs, and temporally resolve the activity of ARX and PAX4 during endocrine lineage segregation due to the presence of the fluorescent reporters as a proximation (proxy) for the translated proteins. The generated line did not carry any chromosomal aberration and passed all routine quality controls (Suppl. Fig. 1C-E). Importantly, the ARX/PAX4 reporter iPSC line is pluripotent and can differentiate into all three germ layers (Suppl. Fig. 1F-H).

To study in greater details *PAX4* and *ARX* expression during β cell differentiation, we differentiated the ARX/PAX4 reporter iPSC line using an adapted version of a previously published β cell differentiation protocol in 3D culture^17^ (Fig. 1D). We monitored the presence and reporter activity of mCherry and CFP as a proxy of *PAX4* and *ARX* expression, respectively, using live flow cytometry and immunofluorescence (IF) (Fig. 1E-F). Cell clusters at early PP stage (S3) showed less than 1% positive cells for both mCherry and CFP by live flow cytometry (Fig. 1E). The percentage of fluorophore-positive cells started to increase in the transition between late PPs and EPs (day 3 of S4 – S4d3) (Suppl. Fig. 2A). While the relative number of cells positive for mCherry peaked at the EP stage (S5d7), the relative number of CFP-positive cells peaked at the SC-β cell stage (S6d14) (Fig. 1E-F; Suppl. Fig. 2A). Since fluorescent reporters are transcribed and translated together and at the same rate as the TFs themselves, presence of the reporters is a proxy for TF activity. Therefore, this data suggests that *PAX4* and *ARX* expression patterns, and therefore their activity, are asynchronous. To test this hypothesis, we stained wild-type clusters (XM001^39^) for the endogenous PAX4 and ARX proteins. We analyzed diverse time points during the differentiation protocol toward SC-β cells and we found almost identical dynamic expression between the endogenous proteins and their fluorescent reporters (Fig. 1F-G; Suppl. Fig. 2). Although onset of fluorescent reporter activity and TF protein production is identical, mCherry seemed to be more stable and detectable for a longer time-span compared to the endogenous PAX4 proteins. While endogenous PAX4 levels decreased starting around S5d6 (Fig. 1F-G; Suppl. Fig 2B-C), the relative numbers of cells positive for mCherry were still elevated at S5d7 (Fig. 1E-F; Suppl. Fig. 2A, 2C), making mCherry a transient lineage tracer for PAX4 in the ARX/PAX4 reporter cell line (Suppl. Fig 2C). Due to the integration of the reporter into histone H2B, the same should be true for ARX-CFP. Finally, onset of *ARX* and *PAX4* expression and their protein synthesis are accurately reflected by our double reporter system which allows us to dynamically resolve temporal changes of the two lineage determining TFs and, consequently, their downstream GRNs. Together with the human fetal expression patterns, this data suggests a sequential expression of *PAX4* and *ARX* during human β cell differentiation in which *PAX4* is transiently co-expressed with *NEUROG3* and transcriptionally active in early- and mid-EPs, while *ARX* is expressed and active in α cell progenitors at later stages of differentiation. If true, this hypothesis would support a different model for human EP differentiation towards the α or β cell fate than proposed *in vivo* in the mouse (Fig. 1A).

### Single-cell multiomic analysis suggests ARX repression of *PAX4* during *in vitro* human SC-β cell differentiation

To test our hypothesis and characterize the regulatory program governing α *versus* β cell fate acquisition during differentiation of human SC-islets, we took advantage of the ARX/PAX4 reporter iPSC line. While S5d4 is one of the first time point in which the dynamics of PAX4-mCherry^+^ and ARX-CFP^+^ cells diverge, clusters at this stage should be enriched in cells going through the transition from late PPs towards EPs (Fig 1D-E), an ideal differentiation time point to study the dichotomy between ARX- and PAX4-mediated lineage determination. We therefore used fluorescence activated cell sorting (FACS) to isolate PAX4-mCherry^+^, ARX-CFP^+^ and PAX4-mCherry^+^/ARX-CFP^+^ cells at S5d4 and performed single cell multiomic analysis of all three fractions together with an unenriched sample (Fig. 2A; Suppl. Fig. 3A). To simultaneously monitor gene expression and chromatin accessibility in single cells, we conducted paired single-nucleus RNA-seq and single-nucleus assay for transposase-accessible chromatin using sequencing (snATAC-seq) resulting in a total of 18’544 cells. We first performed cluster analysis of the two datasets separately, and then, to ensure more precise identification of cell types^40,41^, we integrated both data sets (Suppl. Fig. 3B). This identified nineteen distinct cell clusters (Fig. 2B). One cluster was comprised of non-endocrine cells and one of pre-endocrine cells. Three EP populations were annotated: very early progenitors, early progenitors, and EPs. Four α cell clusters were present: early and late α progenitors, α cells with low expression of glucagon, and α cells. In addition, four β cell clusters were annotated: β progenitors, β cells expressing the neuronal marker *Growth Associated Protein 43* (*GAP43*), β cells expressing both *GAP43* and *secretin* (*SCT*), and β cells. Our data set also contained ε progenitor and ε cell clusters, together with other endocrine cells characterized by the expression of other hormones and polyhormonal cells (Fig. 2B; Suppl. Fig. 3C). We then looked at the relative frequency of cell types across our FACS-sorted fractions (Fig. 2C-D; Suppl. Fig. 3D). As expected, half of the control unsorted sample was comprised of non-endocrine cells, while the other half of EPs and endocrine cells. The PAX4-mCherry^+^ fraction contained mostly EPs, β cell progenitors, and β cells. The ARX-CFP^+^ fraction contained α cells, endocrine cells expressing the hormone *cholecystokinin* (*CCK*), polyhormonal cells, α cell progenitors, and ε cells, which were previously hardly detected in any other single cell datasets^18,38,40^. On the other hand, and similarly to the ARX-CFP^+^ sample, the fraction enriched for double positive cells for both ARX and PAX4 reporters (ARX-CFP^+^/PAX4-mCherry^+^) was constituted mainly by α cell progenitors, α cells, polyhormonal and *CCK*-expressing cells (Fig. 2C-D; Suppl. fig. 3D). In agreement with our flow cytometry and IF analysis (Fig. 1E-G) and the publicly available scRNA-seq previously analyzed (Suppl. Fig. 1A-B), *PAX4* expression mostly coincided with *NEUROG3* expression in EPs (from very early on to α and β cell progenitors), while *ARX* expression was elevated in α cell progenitors, α cells, polyhormonal, *CCK,* and *GHRL* expressing cells (Fig. 2E; Suppl. Fig. 3E-H). In conclusion, multiomic analysis showed that 1) due to their co-translational cleavage, simultaneous *ARX* and *PAX4* mRNAs and reporter activity reflects TF transcription and activity in our system, 2) *PAX4* expression is transient expressed in EPs, therefore mCherry is a transient *PAX4* reporter in emerging α cell progenitors, and 3) *PAX4* expression, and therefore *mCherry* activity, are maintained in β cell progenitors.

**Fig. 2.**
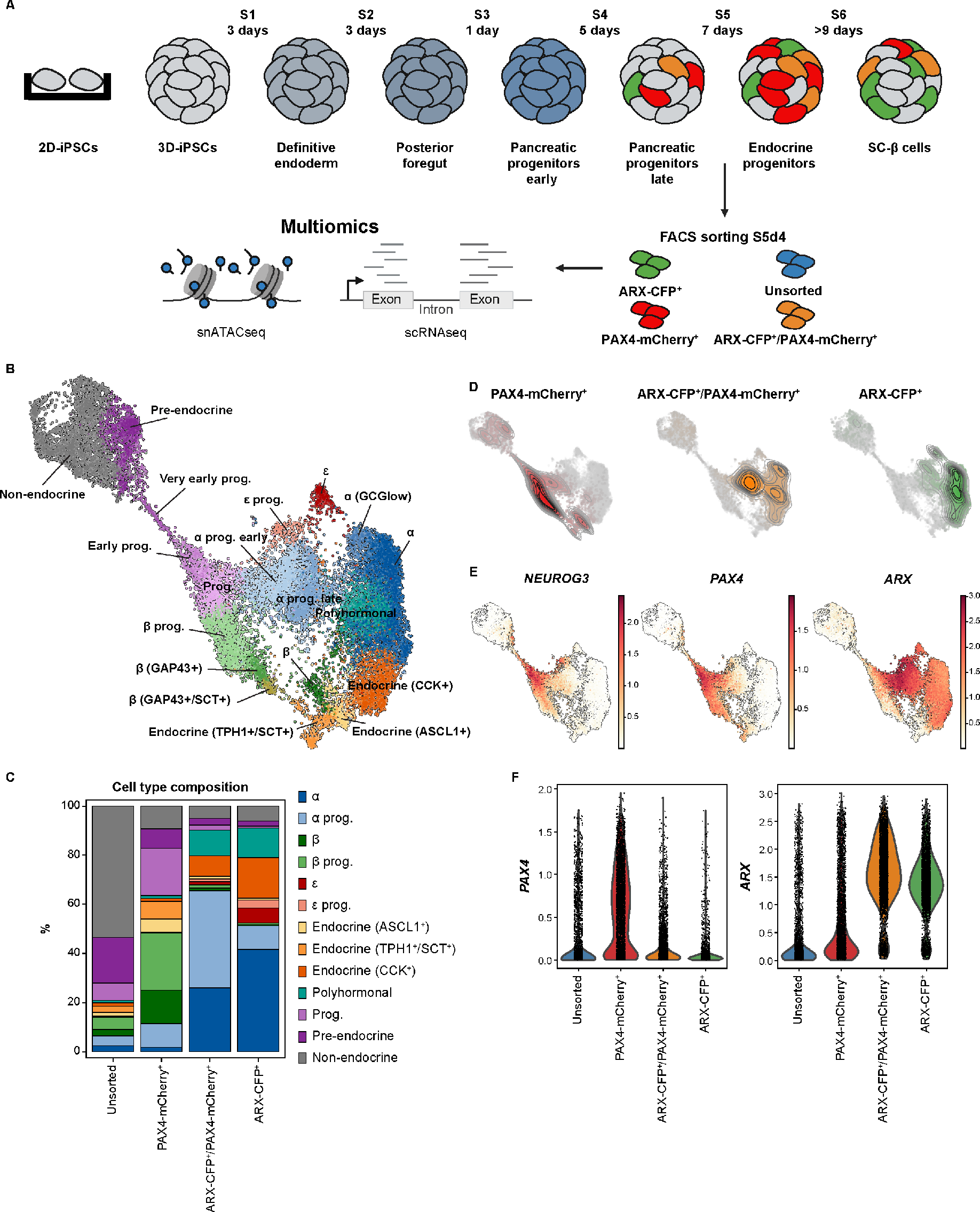
Single cells multiomics suggests ARX repression of PAX4 during in vitro SC-β cell differentiation. **(A)** Experimental strategy used for multiome analysis of ARX-CFP^+^, ARX-CFP^+^/PAX4-mCherry^+^ and PAX4-mCherry^+^ FACS-sorted fractions. Unsorted-live cells were used as control. **(B)** Annotated multiomic UMAP showing the integration of all samples to identify cell clusters. **(C)** Relative frequencies of cell types in each dataset as per annotated UMAP in B. **(D)** Embedding densities of PAX4-mCherry^+^, ARX-CFP^+^/PAX4-mCherry^+^, and ARX-CFP^+^ samples. **(E)** UMAPs showing *NEUROG3* and *PAX4* expression starting from early EPs, and *ARX* expression in late EPs and α-cell progenitors. **(F)** Violin plots showing normalized *PAX4* and *ARX* expression in ARX-CFP^+^, ARX-CFP^+^/PAX4-mCherry^+^, PAX4-mCherry^+^ and unsorted fractions.

To validate our scRNA-Seq data, we performed bulk proteomic analysis of FACS-sorted fractions at S5d4 (Suppl. Fig. 4A). Gene ontology analysis on differentially abundant proteins between unsorted and enriched samples revealed commitment towards β cells of PAX4-mCherry^+^ fractions, and α cells of both the PAX4-mCherry^+^/ARX-CFP^+^ and ARX-CFP^+^ fractions (Suppl. Fig. 4B). In agreement with EPs being postmitotic and less adhesive to the extracellular matrix (ECM)^42–47^, all three samples showed downregulation of proteins related to ECM organization and cell cycle (Suppl. Fig. 4B). Also, the transcriptomic (Fig. 2C-D) and proteomic gene ontology data of the PAX4-mCherry^+^/ARX-CFP^+^ and ARX-CFP^+^ samples showed very similar proteomic profiles (Suppl. Fig. 4C).

In accordance with our hypothesis, these data suggest a sequential expression of *PAX4* and *ARX* during human SC-α and β cell differentiation. In this scenario, the transient action of PAX4 would (directly or indirectly) induce *ARX* expression only in some EPs. Once translated, ARX would be responsible for the direct or indirect repression of *PAX4* mRNA expression in α cell progenitors, promoting α cell differentiation.

### ARX TF action determines α *versus* β cell fate decision

To functionally test this hypothesis, we performed *in vitro* potency assays. We sorted PAX4-mCherry^+^, ARX-CFP^+^, and PAX4-mCherry^+^/ARX-CFP^+^ at EP stage (S5d7) and differentiated them toward SC-β cells separately for two weeks (Fig. 3A-B). As expected, most cells within the PAX4-mCherry^+^-enriched cultures differentiated into insulin-producing SC-β cells (Fig. 3C). On the other hand, most cells within both ARX-CFP^+^- and PAX4-mCherry^+^/ARX-CFP^+^-enriched cultures differentiated into glucagon-producing SC-α cells (Fig. 3C). IF for PAX4 and ARX reporters revealed that, while single positive cells for one reporter (being mCherry for PAX4 or CFP for ARX) kept producing it during the entire culture, PAX4-mCherry^+^/ARX-CFP^+^ cells eventually decreased mCherry production, producing only CFP by the end of the culture (Fig. 3C).

**Fig. 3.**
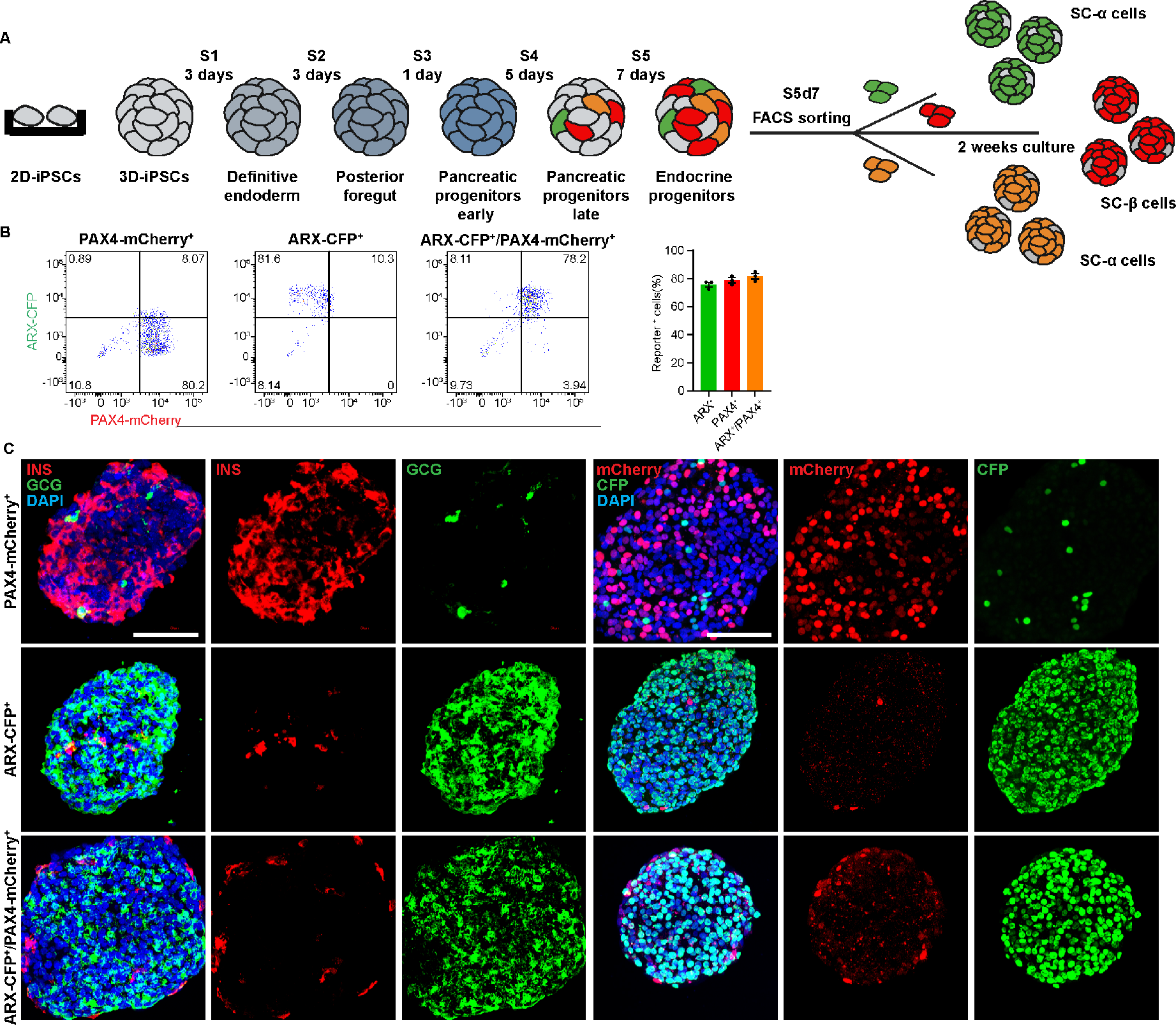
ARX expression coordinates α versus β cell fate decision. **(A)** Experimental strategy for potency assay of ARX-CFP^+^, ARX-CFP^+^/PAX4-mCherry^+^ and PAX4-mCherry^+^ FACS-sorted fractions. **(B)** Representative flow cytometry plots of fractions purity directly after sorting. **(C)** Representative maximum intensity projections of Z-stack confocal acquisitions of FACS-sorted ARX-CFP^+^, ARX-CFP^+^/PAX4-mCherry^+^ and PAX4-mCherry^+^ fractions-derived clusters showing ARX-CFP and PAX4-mCherry, insulin, and glucagon after 2 weeks in culture. Data are presented as mean ± SE. INS: insulin, GCG: glucagon. Scale bars: 50 µm.

Taking together, multiomics, proteomics and potency assays suggested for the first time the lack of stochasticity in the expression of *PAX4 versus ARX* in double positive cells, and highlights ARX expression and activity as a major determinant of α cell fate determination. Consequently, cells expressing both TFs in the mid/late EP stage of endocrinogenesis commit to the α cell lineage.

### ARX and PAX4 participate in a GRN amenable for manipulation

To study if and how ARX and PAX4 modulate each other’s molecular and cellular activity, we queried our single cell ATAC-Seq data for potential PAX4 and ARX binding sites within each other’s putative cis-regulatory elements (CRE). While we found motifs of each TF in putative CREs of the other gene, there was no correlation between the accessibility of the CRE and TF expression. We thus concluded that there are no binding sites of PAX4 within the *ARX locus* and *vice versa* (Fig. 4A). We instead found a putative ARX binding site within the open chromatin of its own *locus*, suggesting a possible feedback loop by which ARX could promote its own transcription (Fig. 4A). We then used the integration of our scRNA-Seq and snATAC-Seq datasets to compute a GRN involving *PAX4* and *ARX* during α and β cell differentiation (Fig. 4B). From the nature of the correlations found within this network, ARX seemed to act more often as a transcriptional repressor, while PAX4 seemed to be mostly acting as a transcriptional activator. The GRN suggested furthermore that 1) ARX can regulate itself, and 2) ARX and PAX4 do not directly interact but 3) diverse intermediate TFs are involved in *ARX* and *PAX4* transcriptional regulation.

**Fig. 4.**
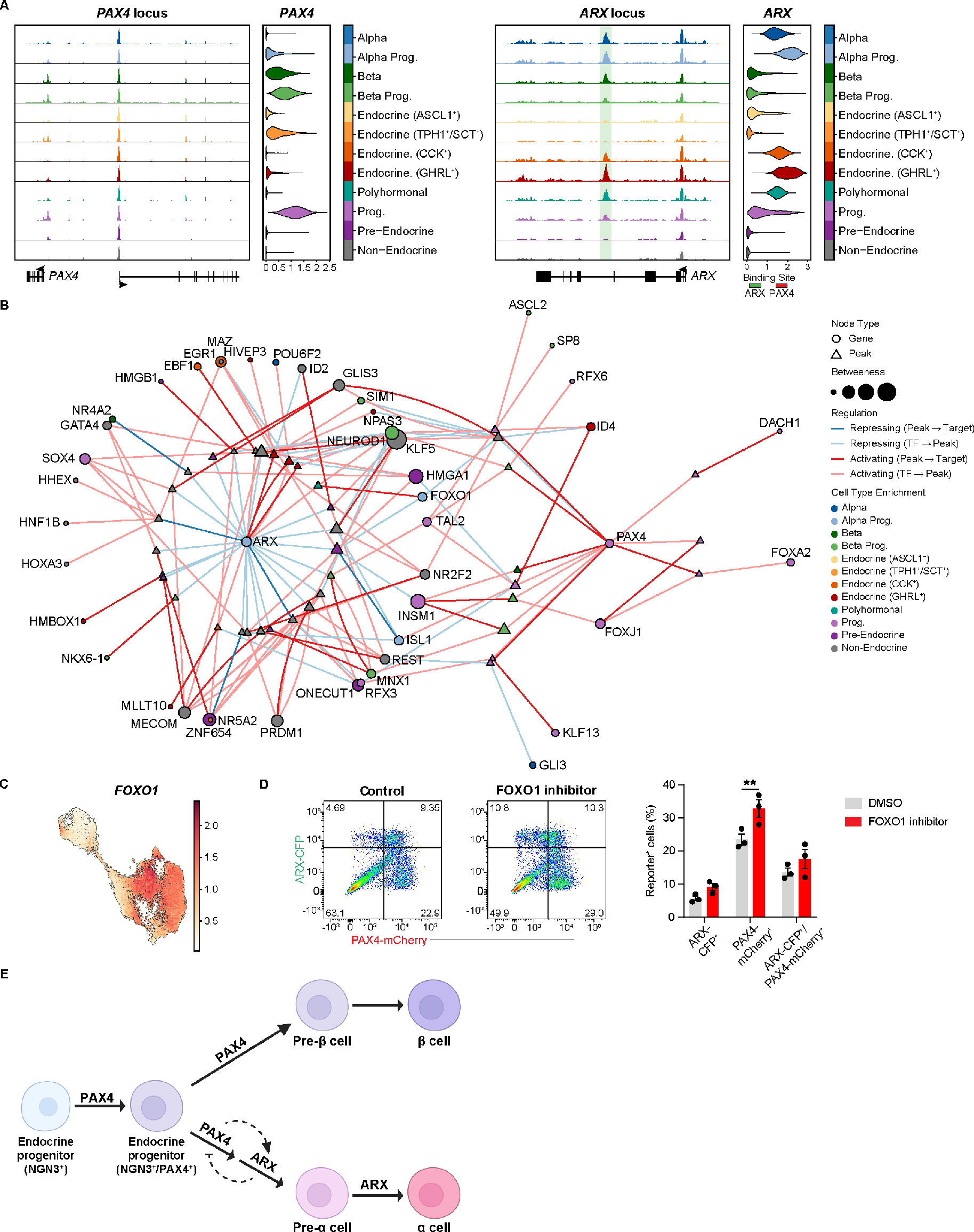
ARX and PAX4 participate in a GRN that can be manipulated. **(A)** ATAC sequencing peaks showing chromatin accessibility within the *PAX4* and *ARX* CREs together with *in-silico* predicted PAX4 and ARX binding sites. Note how ARX could only bind its own locus (green shaded putative binding site) but not the *PAX4* locus. **(B)** Prediction of a GRN involving PAX4 and ARX using multiomic datasets from ARX-CFP^+^, ARX-CFP^+^/PAX4-mCherry^+^ and PAX4-mCherry^+^ FACS-sorted fractions. **(C)** UMAPs showing enriched *FOXO1* expression in α-cell progenitors. **(D)** Representative flow cytometry plot (left) and related quantification (right) of the percentage of ARX-CFP^+^, ARX-CFP^+^/PAX4-mCherry^+^ and PAX4-mCherry^+^ cells at S5d7 (EPs) after treatment with FOXO1 inhibitor starting from S5d1. **(E)** Working model of ARX and PAX4 mechanism of action during human *in vitro* α *versus* β cell fate decision per our data. Dashed lines represent indirect transcriptional activation of *ARX* by PAX4, and indirect repression of *PAX4* transcription by ARX. Data are presented as mean ± SE. Two-ways ANOVA with Šídák multiple comparison test. ** ≤ 0.01.

In addition to TFs upstream of *ARX* and *PAX4*, our GRN allows for identification of their putative downstream targets. For example, ARX seems to activate *FOXO1* transcription in α progenitor cells (Fig. 4B-C). In both murine and human pancreas, *FOXO1* expression is present in all pancreatic cells during development to then become restricted to β cells postnatally^48,49^. In the adult islets, FOXO1 regulates β cell proliferation and mediates their response to oxidative stress, while *FOXO1* knock-down or inhibition results in dedifferentiation and loss of cell identity^31,50–52^. To test the validity of our GRN and our ability to manipulate it, we treated emerging ARX/PAX4 reporter SC-islets with the small molecule inhibitor of FOXO1 (AS1842856), which has already been shown to be able to improve human embryonic stem cell (hESC) differentiation towards β cells *in vitro*^53^. FOXO1 inhibition during the entire S5 resulted in a significant increase (10%) in the number of PAX4-mCherry^+^ cells at S5d7, compared to cells treated with DMSO vehicle alone (Fig. 4D). Thus, our GRN analysis using the ARX/PAX4 double reporter line identifies druggable targets to manipulate and study human α *versus* β cell fate decision *in vitro*.

Taken together these results suggest a model of human β cells differentiation in which PAX4 and ARX do not stochastically cross inhibit each other, but instead PAX4 in EPs indirectly induces *ARX* expression, that in turn indirectly represses *PAX4* expression, inducing differentiation toward the α cell lineage (Fig. 4E).

### Artemether has a profound effect on endocrine induction lineage induction and segregation

The generation of the double ARX/PAX4 reporter iPSC line opens new avenues to study simultaneously lineage trajectories on temporal resolved single cell level and the more general biological consequences of small molecule treatments on α *versus* β cell fate decision during SC-islet differentiation. Here we tested the use of the FDA-approved antimalaria drug artemether, which has been reported to trigger α-to-β transdifferentiation in some but not all studies^30–33,35^. Therefore, we continuously treated ARX/PAX4 differentiating reporter cells starting from early PP until SC-β cell (S4 to end of S5) stage and performed multiomics analysis (scRNA-seq and snATAC-seq) on unsorted clusters at S5d4 and live flow cytometry at S5d7 (Fig. 5A; Suppl. Fig 5B). Multiomic analysis showed increased endocrine, β, and α progenitor cell numbers at the expenses of both non-endocrine and more differentiated cell types upon artemether treatment (Fig. 5B-C). Accordingly, differential gene expression analysis of each untreated cluster *versus* its artemether-treated counterpart revealed increased expression of EP markers (such as *NEUROG3* and *INSM1*) upon treatment, while hormone expression levels were reduced (Suppl. Fig. 5A). Together, this data suggests a remarkable increase in endocrine induction and slower differentiation of artemether-treated cells. Confirming this hypothesis, flow cytometry analysis three days later (S5d7) revealed a significantly increased percentage of PAX4-mCherry^+^ cells upon artemether treatment, suggesting an enrichment for both endocrine and β cell progenitors (Fig. 5D). We confirmed this hypothesis with flow cytometry and IF analysis, which also revealed increased number of cells positive for EP markers (NEUROG3, NKX2.2, NKX6.1) and insulin, with a reduction in the percentage of glucagon-positive cells and bihormonal cells (insulin^+^/glucagon^+^) (Fig. 5E-F). Continuous treatment with artemether until the end of the culture at S6d14 resulted in a strong reduction of glucagon-positive cells and a trend towards increased number of insulin-positive cells, as shown by flow cytometry (Fig 5G; Suppl. Fig. 5C), suggesting that artemether treatment has an impact on α *versus* β cell fate decision.

**Fig. 5.**
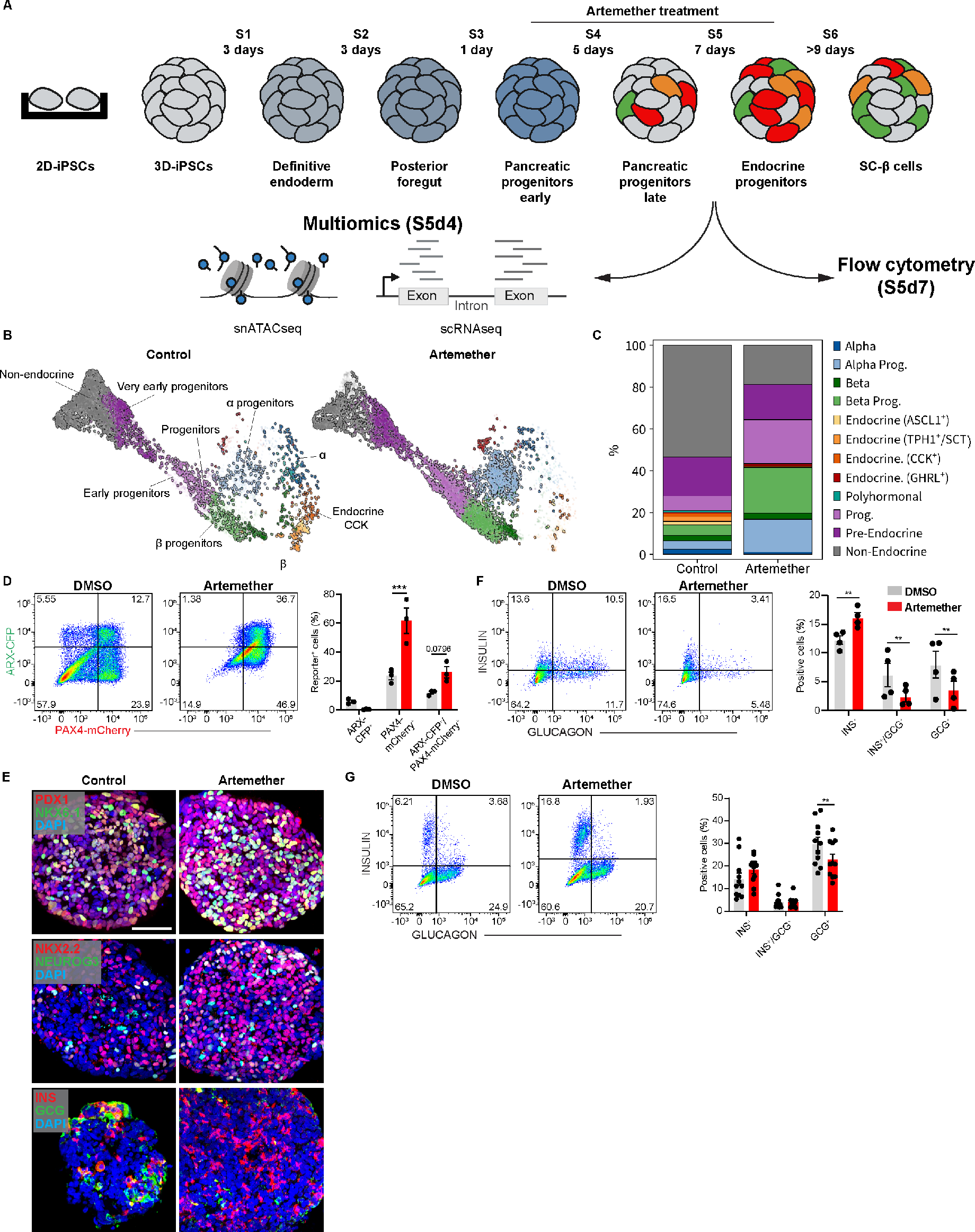
Artemether has a profound effect on endocrine lineage induction and segregation. **(A)** Experimental strategy for artemether treatment followed by multiomic analysis at S5d4 and flow cytometry analysis at S5d7. **(B-C)** Multiome UMAPs of unsorted (4609 cells) and artemether-treated samples (4607 cells) (B) and relative cell type composition (C). **(D)** Representative flow cytometry plots and related quantification of the percentage of ARX-CFP^+^, ARX-CFP^+^/PAX4-mCherry^+^ and PAX4-mCherry^+^ cells at S5d7 after artemether treatment. **(E)** Representative maximum intensity projections of Z-stack confocal acquisitions of IF staining showing EP markers (PDX1, NKX6.1, NKX2.2, and NEUROG3), insulin and glucagon hormones within artemether-treated and control clusters at S5d7. **(F-G)** Representative flow cytometry plots and related quantification of hormone-positive cells at S5d7 (F) and S6d14 (G) after continuous treatment with artemether starting from S4. Data are presented as mean ± SE. Two-ways ANOVA with Šídák multiple comparison test. ** ≤ 0.01; *** ≤ 0.001. INS: insulin, GCG: glucagon. Scale bars: 50 µm.

### Artemether induces cell proliferation while protecting against cell death

In addition to the altered composition of SC-islets, artemether-treated clusters were bigger than DMSO-treated controls (Fig. 6A). To understand what drives this phenotype, we quantified cell proliferation and death using flow cytometry. The percentage of proliferating cells (Ki67-positive) was increased specifically at S4d5 controls (Fig. 5B-C), whereas the percentage of apoptotic cells (cleaved Caspase3 [clCasp3]-positive) was decreased at the EP stage in artemether-treated samples compared to DMSO-treated (Fig. 6D-E). Together, we showed how artemether treatment could improve SC-β cell differentiation protocols: it promotes PP expansion, increases EP induction and positively affects SC-islets cellular composition, making artemether a potential drug to direct SC-islet differentiation.

**Fig. 6.**
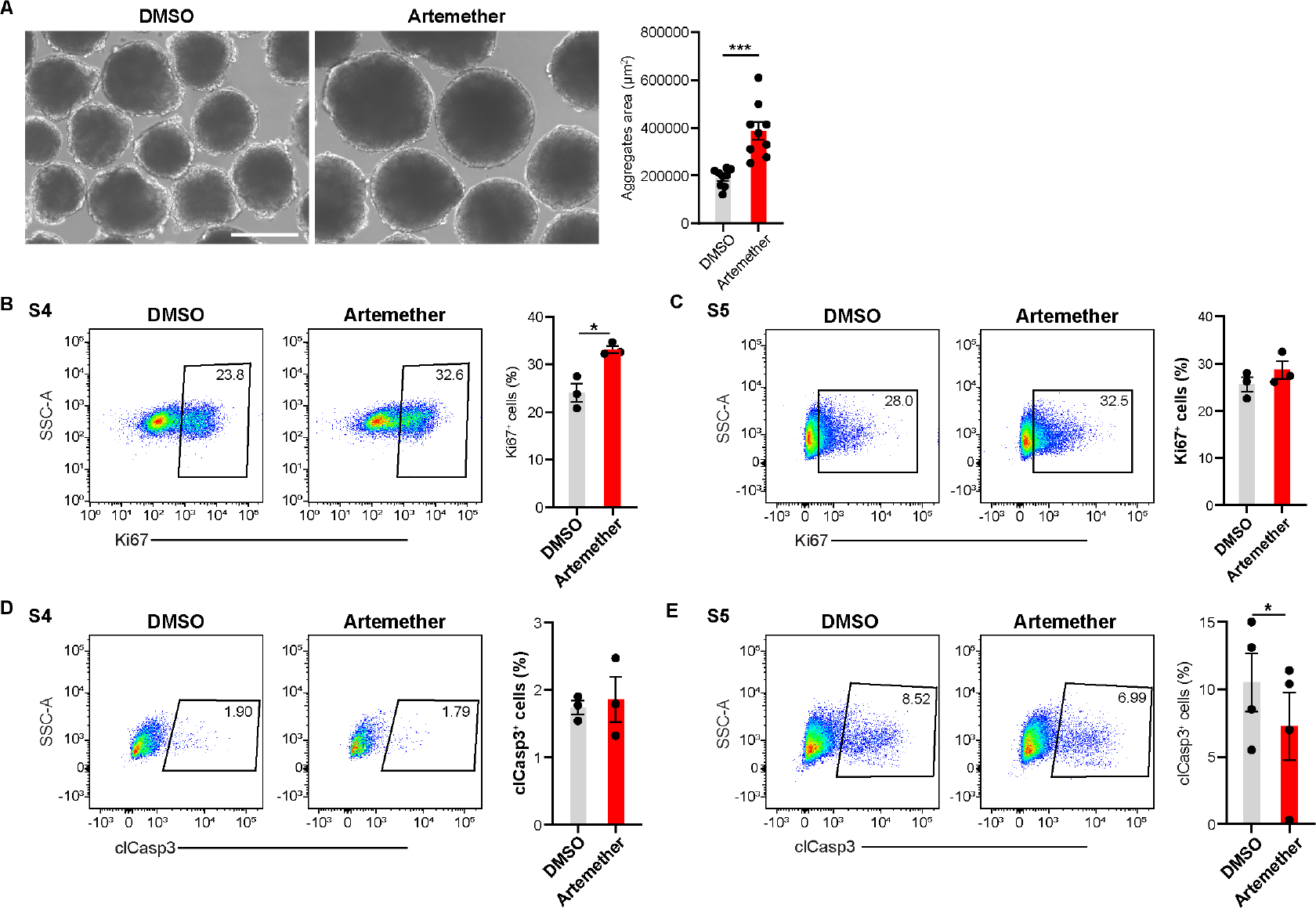
Artemether induces cell proliferation while protecting against cell death. **(A)** Representative bright field pictures and related area quantification of clusters treated with artemether or DMSO at S5d7. Note the size increase and the increased sphericity of the artemether-treated clusters. **(B-C)** Representative flow cytometry plots and related quantifications of the percentage of Ki67-positive cells at S4d5 (B) and S5d7 (C), after DMSO or artemether treatment. **(D-E)** Representative flow cytometry plots and related quantifications of the percentage of clCasp3-positive cells at S4d5 (D) and S5d7 (E), after DMSO or artemether treatment. Data are presented as mean ± SE. Two-ways ANOVA with Šídák multiple comparison test. * ≤ 0.05. Scale bar: 300 µm.

## DISCUSSION

The role of ARX and PAX4 during human pancreas endocrinogenesis has not been fully understood, their upstream and downstream regulation has not been characterized and the dichotomy between the two TFs has never been fully demonstrated. Here, we generated a human iPSC double reporter line allowing to test the dogma by which the activity of the two TFs ARX and PAX4 in EPs would stochastically drive α or β cell fate decision, respectively. Following multiomics analysis we generated a GRN centered around ARX and PAX4 and propose a new model of human α *versus* β cell fate decision. We showcased the ARX/PAX4 reporter line both as a tool to dissect genetic regulation underlying human endocrinogenesis, and as a new platform for drug discovery uncovering its potential to improve SC-islet differentiation protocols. We showed artemether-driven EP induction and decreased α cell differentiation, together with PP expansion, suggesting artemether as a potential new chemical to improve the yield of PPs and EPs during direct SC-β cell differentiation. On the other hand, the effect of artemether treatment on *ARX* and *PAX4* transcription seems negligible in our system.

Our study showed that *ARX* and *PAX4* temporal and spatial expression patterns do not largely coincide during human endocrinogenesis. We proved that both *in vitro* and *in vivo PAX4* is transiently expressed in EPs slightly after *NEUROG3*, while *ARX* is mostly expressed in cells already committed to the α cell lineage. This data is in line with recent human first trimester fetal single cell multiomics data, highlighting conserved and divergent transcriptional regulations during human and murine endocrinogenesis^54^. Consequently, our work suggests *PAX4* as a transient EP marker, downstream of *NEUROG3*. Our new model of human endocrinogenesis does not fully resemble the one from Ma and colleagues^54^, as it does not include an *ARX*-positive ε cell differentiation branch. This might be due to temporal discrepancies between SC-islets differentiation and *in vivo* endocrinogenesis. In line with the presence of this early branch of differentiation that establishes ε cell commitment before α and β cell fate decision, samples embedding within our multiomics analysis showed contribution to the *ε* progenitors and *ε* cell clusters only from the ARX-CFP^+^ sample but not from the ARX-CFP^+^/PAX4-mCherry^+^ (Fig. 2D). While we did not find a direct connection between *NEUROG3* and *PAX4*, our GRN showed both *Neuronal Differentiation 1* (*NEUROD1*) and *Insulinoma-associated 1* (*INSM1*) as direct targets of *NEUROG3.* Both TFs would, in turn, activate *PAX4* transcription. This data are in agreement with mouse studies showing how both *NEUROD1* and *INSM1* reinforce endocrine cell fate acquisition downstream of *NEUROG3*^55–61^. In addition, while PAX4 has been classically described as a repressor of hormone transcription^62–64^, it acts mainly as an activator at the EP stage in our system. On the other hand, ARX exhibits mainly a transcriptional repressor activity, with the notable exception of the feedback loop promoting its own transcription (Fig. 4B).

We showed how useful the new ARX/PAX4 double reporter line could be as a drug screening platform. In combination with our GRN, it allows identification and testing of druggable hubs for the optimization of differentiation protocols, with the goal of generating SC-islets with proportions of α and β cells similar to human islets. It can also be used to simultaneously test the mechanisms of action and impact on SC-islet differentiation of putative α-to-β transdifferentiation inducers in a relevant human model system. Premature *FOXO1* activation in murine pancreatic progenitors *in vivo* was shown to increase α cells, while its specific KO in pancreatic progenitors led to increase β cell numbers^49^. Other studies using animal models and mature islets identified FOXO1 inhibition as a potent dedifferentiation cue^50,65,66^. The results of FOXO1 inhibition in our system are in line with previous literature where FOXO1 inhibition was shown to improve hESC differentiation towards β cells *in vitro* and reduce *ARX* expression in mature murine and human islets^31,53^. On the other hand, the consequences of treatment with artemisinin and its derivatives on pancreatic cell fate decision and regeneration have been controversial. Artemether was reported to modulate endocrine cell fate commitment and promote β cells regeneration in immortalized α cell lines, murine and human primary islets, *Danio Rerio*, and mice^30–32^. Others studies reported instead dedifferentiation of murine and human primary islets ^33^ or no changes upon treatment *in vivo* in mice^35,67^. While none of these studies looked at artemether potential during human endocrinogenesis, in this context our data provide strong evidence that artemether induces endocrine induction and favors β cells at the expenses of α cell differentiation.

Finally, our study provides novel and detailed insights into the regulation of human α *versus* β cell fate decision both *in vitro* and *in vivo* that could pave the way to the establishment of new and better protocol for the differentiation of SC-islets for cell replacement therapy, as well as a new human-relevant drug testing platform for regenerative therapy purposes.

## METHODS

## 1. Generiation of hiPSC line

### 1.1. Cloning of targeting constructs

HMGUi001-A-46 (a.k.a. ARX^nCFP/nCFP^; PAX4^mCherry/mCherry^) transgenic double-reporter hiPSC line was generated by insertion of mCherry in the PAX4 allele of HMGUi001-A-4 hiPSC line^36^. First, PAX4-single guide RNA was cloned into the pu6-(Bbsl) sgRNA-CAG-Cas9-Venus-bpA plasmid (Addgene plasmid #86986) via Gibson assembly which introduced a double-strand break at the N-terminus of Exon 1, specifically after 10 bp from the start codon. Double-strand break was repaired by a targeting vector of Histone 2B-mCherry-RGSHis-T2A construct flanked by a 703 bp 5’ homology arm (5’HA) and a 1008 bp 3’ homology arm (3’HA). Both homology arms were amplified from genomic DNA of HMGUi001-A hiPSCs (a.k.a. XM001) by PCR.

### 1.2. Transfection

hiPSC transfection was performed according to previous studies^68^. Briefly, HMGUi001-A-4 hiPSCs were seeded at a density of 0.4×10^6^ cells/well of 6 well tissue culture plate with StemMACS™ iPS-Brew XF medium (Miltenyi Biotec cat#130-104-368) and 10 µM Y-27632 (SantaCruz cat# sc-281642A). Next day, the medium was replaced without Y-27632 for at least 4 hours before proceeding with the transfection. The targeting construct, H2B-mCherry-RGSHis-2A and targeting plasmid, pu6- (Bbsl)PAX4-sgRNA-CAG-Cas9-Venus-bpA were delivered via Lipofectamine™ Stem Transfection Reagent (Fisher Scientific, Cat# STEM00003). 48 hours after transfection, cells were treated with EDTA for 5 minutes and then harvested with StemMACS™ iPS-Brew XF supplemented with 10 µM Y-27632. Cell suspension was collected and centrifuged at 1400 rpm for 4 minutes and then pellet was resuspended and filtered through low attachment polypropylene flow cytometry tubes. Transfection efficiencies were assessed by recording the Venus-positive cells via flow cytometry. 3000-4000 Venus-positive cells were sorted and seeded onto pre-filled Geltrex (Thermo Fisher Scientific, Gibco cat#A1413302)-coated 10cm tissue culture plate (Thermo Fisher Scientific, cat#150350).

## 2. hiPSC Culture

### 2.1 Adherent culture

HMGUi001-A-46 hiPSCs were cultured on tissue culture plate coated with 1:100 Geltrex in StemMACS™ iPSBrew XF culture medium at a density of 1×10^5^ cells/cm^2^ under standard culture conditions (37°C, 5% CO2, and 95% humidity). The medium was replaced daily, and cells were passaged every 3-4 days once 70% confluency was reached. Culture were rinsed with Dulbecco phosphate-buffered saline (DPBS, no Calcium, no Magnesium, Thermo Fisher Scientific, Gibco cat#14190094) and incubated with Accutase (Merck, Sigma-Aldrich cat# A6964-100ml) for 4-5 minutes at 37°C. Dispersed hiPSCs were spun at 1400 rpm for 4 minutes at room temperature. hiPSCs were resuspended in StemMACS™ iPS-Brew XF medium with 10 µM Y-27632 and seeded on pre-filled Geltrex-coated dishes. After 24 hours, the medium was replaced without Y-27632.

### 2.2 Suspension culture

HMGUi001-A-46 hiPSCs were dispersed using Accutase (Merck, Sigma-Aldrich cat# A6964-100ml) and counted using counting chambers (Hirschmann EM techcolor cat#8100204) 1×10^6^ cells/ml were seeded into prefilled 30 ml spinner flask with StemMACS™ iPS-Brew XF medium with 10 µM Y-27632 and placed onto magnetic stirrer platform (ABLE Biott cat# ABBWBP03N0S-6) at 60 rpm. Clusters of approximately 100 µm in diameter formed within 48 hours. The clusters were splitted every 3 day once a diameter of 200-300 um was reached. They were collected and rinsed with DPBS and incubated with Accutase for 7 minutes at 37°C. The clusters were gently dispersed by pipetting and filtered using 100 µm cell strainer (Corning, cat# 431752). 1×10^6^ cells/ml cells were transferred into prefilled 30 ml spinner flask with StemMACS™ iPS-Brew XF medium with 10 µM Y-27632. The medium was replaced without Y-27632 within 48 hours.

### 2.3 hiPSCs differentiation

HMGUi001-A-46 line was differentiated towards SC-β cells following the protocol outlined by Velazco-Cruz and colleagues ^17^ after 2-3 passages in suspension culture. hiPSCs were seeded at a density of 1×10^6^ cells/ml in a 30 ml spinner flask for differentiation. Subsequently, differentiating clusters were transferred to 6-well Ultra-Low binding plates (Corning, Falcon cat#1015443) at a density of 5×10^6^ cells/well at the beginning of the S4 or S5 for further treatments.

## 3. Live flow cytometry and sorting

For live flow cytometry, 30 differentiating clusters were incubated with Accutase for 5 minutes in a water bath and dispersed to single cells. mCherry and CFP reporters were analyzed through PE-Texas Red and CFP channels, respectively, at S3-S6 using a FACS Aria III (Becton Dickinson) operated with BD FACSDiva software. A total of 30000 events was recorded. FlowJo software (Becton Dickinson, version 10.7.1) was used for downstream analysis. For the sorting, clusters at S5 days 4 and 7 were dispersed into single cells and enriched based on CFP and mCherry fluorescence. The sorting parameters were established in FACS DIVA software. The clusters were dispersed using Accutase, then 10×10^6^ cells were resuspended in 2 ml of StemMACS™ iPS-Brew XF supplemented with 10 µM Y-27632 and transferred into polystyrene round bottom FACS tube (Corning, Falcon cat#3522235). PAX4-mCherry^+^, ARX-CFP^+^/PAX4-mCherry^+^, and ARX-CFP^+^ cells were then sorted into polypropylene round bottom FACS tube (Corning, Falcon cat#352063). Sorting accuracy was verified by re-analyzing 100 µl of each sorted population.

## 4. Flow cytometry staining

Clusters were dispersed into single cells at indicated differentiation stages. Cells were blocked and permeabilized using Donkey Block (DB - PBS 1x, 10% FBS, heat inactivated, 0.1% Tween-20, 0.1% BSA, 3% Donkey serum) buffer containing 0.2% Triton X-100 (Merck, Sigma-Aldrich cat# T8787-250 ML) for one hour at room temperature. The cells were then incubated with primary antibodies diluted in DB+0.2% Triton X-100 overnight at 4°C. Next day, the cells were spun down, washed twice with DPBS, and incubated with secondary antibodies for two hours at room temperature. After staining, the cells were washed three times with DPBS and processed through flow cytometry. FACS Diva Software was used to plot the data, matching the specific fluorescence channels of the antibodies used. Controls included stained HMGUi001-A-46 hiPSCs and samples stained only with secondary antibodies. All experiments were analyzed using FlowJo software (Becton Dickinson, version 10.7.1). Lists of the used primary and secondary antibodies can be found in Materials table 1-2.

## 5. Clusters sectioning and staining

50 clusters were collected, washed with DPBS, fixed with ice-cold 4% Paraformaldehyde (PFA; Boster cat # BSBTAR1068) for 20 minutes at room temperature. They were then immersed in a 10% (w/vol) D-(+)-Sucrose (ITW Reagents, PanReac AppliChem cat#57-50-1) solution for 2 hours at room temperature, followed by a 30% (w/vol) sucrose and incubated overnight in a 1:1 mixture of 30% sucrose and Tissue Freezing Medium (Leica, (14)020108926) in 4°C. The next day, the clusters were embedded in cryomolds (Tissue-Tek cat#4566) filled with Tissue Freezing Medium, frozen on dry ice for 1 hour, and then transferred to −80°C. The clusters were sectioned at 15 μm thickness using a Cryostat (Leica, CM1860). For staining, the sections were rehydrated in DPBS for 30 minutes and permeabilized in DPBS with 0.2% Triton X-100 and 1M Glycin (Merck, Sigma-Aldrich cat#G8898-1KG). The slides were blocked with DB for 2 hours at room temperature and then incubated overnight at 4°C with primary antibodies diluted in DB. Next day the slides were washed in DPBS, incubated with secondary antibodies diluted in DB for 2 hours at room temperature, and stained using DAPI (2 µg/mL) for 20 minutes at room temperature. Finally, slides were mounted using Elvanol (25% Glycerol, 10% Mowiol, 100mM Tris pH 8.0, 2% DABCO in ddH_2_O) and left to dry overnight at room temperature. Imaging was performed on a Zeiss LSM 700 at 63x magnification and 1x zoom, followed by image processing and analysis using ImageJ and Photoshop. Zen Blue software was use to generate maximum projections of Z-stack acquisitions.

## 6. hiPSCs characterization

### 6.1 hiPSCs pluripotency analysis

Adherent HMGUi001-A-46 hiPSCs were dissociated using Accutase for 5 minutes at 37°C, spun down at 1400 rpm for 4 minutes, and resuspended in StemMACS™ iPS-Brew XF medium with 10 µM Y-27632. 1×10^6^ cells were stained with SSEA-4-488-FITC and TRA-1-60-PE FACS antibodies. The control samples were stained with isotype control (REA-Control)-FACS antibodies (Miltenyi Biotec cat# 130-104-610, 130-107-146) according to the manufacturer’s instructions (Miltenlyi Biotec). The samples were analyzed using FACS Aria III, 30000 events per sample were recorded.

### 6.2 Three germ layer differentiation

HMGUi001-A-46 hiPSCs were differentiated towards endoderm, ectoderm and mesoderm using the StemMACS™ Trilineage Differentiation Kit (Miltenyi Biotec, Cat# 130-115-660) according to manufacturer’s instructions.

### 6.3 Short tandem repeat polymerase chain reaction analysis

STR analysis was performed by Genomic Core Facility, Helmholtz Zentrum München. Briefly, STR analysis was performed with the AmpFLSTR Identifiler PCR Amplification Kit (Thermo Fisher Scientific, Darmstadt, Germany) using 10 ng of genomic DNA according to the manufacturer’s recommendation. This kit amplifies 15 STR markers and amelogenin in one multiplex assay. Amplification products were mixed with GeneScan 500 LIZ size standard (Thermo Fisher Scientific, Darmstadt, Germany) and separated by size using capillary electrophoresis on a SeqStudio Genetic Analyzer (Thermo Fisher Scientific, Darmstadt, Germany). Data analysis was performed with GeneMapper 5.0 software (Thermo Fisher Scientific, Darmstadt, Germany).

### 6.4 Karyotyping

Karyotyping was performed by Institute of Human Genetics, Technische Universit at München. Briefly, 10-30% confluent iPSC were treated with colcemid to inhibit cell division, trypsinised, hypotonised, and fixed. Finally, the cell suspension was applied to several slides, dried overnight, and stained with Giemsa stain. At least 20 metaphases were microscopically acquired and analyzed using the Applied Spectral Imaging software.

## 7. Single-cell sequencing and data analysis

### 7.1. Single cell suspension preparation

The clusters were dissociated at S5 day 4 into single cells. 1×10^5^ PAX4-mCherry^+^, ARX-CFP^+^/PAX4-mCherry^+^, and ARX-CFP^+^ cells were sorted as described in section 3. Additionally, 1×10^5^ unsorted live cells were enriched using the DAPI channel, serving as control for multiomics.

### 7.2. Multiome (scRNA/snATAC)

For nuclei isolation and library construction, a low input nuclei isolation protocol adapted from 10X Genomics was performed. In brief, sorted cells were washed once with 1 mL PBS + 1% BSA, counted, centrifuged, and supernatant was aspirated. Subsequently, the washed cell pellet was resuspended in chilled lysis buffer with 0.5x detergent concentration (50 μL per sample) and placed on ice for 5 min. Then wash buffer (500 μL per sample) was added and nuclei were centrifuged. To gradually change from wash to diluted nuclei buffer, cells were washed once in a 1:1 mixture of wash buffer and diluted nuclei buffer and subsequently one with pure diluted nuclei buffer. The washed isolated nuclei were then resuspended in 7-10 μL diluted nuclei buffer and were directly added to the transposition reaction after quality control and counting. In all following steps, 10X Genomics’ Single Cell Multiome ATAC and gene-expression protocols were followed according to the manufacturer’s specifications and guidelines. The final libraries were sequenced on the Illumina NovaSeq 6000 platform following the recommendations from 10X Genomics. Raw reads were aligned to the improved pig genome annotation and pre-processed using the 10X Genomics CellRangerARC pipeline (v 2.0.0) for downstream analyses.

### 7.3. Preprocessing of 10X multiome raw data

Multiome data was preprocessed using Scanpy^69^ (v1.9.1) and Muon^70^ v0.1.3). *Filtering of low-quality cells.* First, DropletUtils^71^ (v1.14.2) was used with default parameters to estimate ambient gene expression probabilities. Next, each sample was assessed seperately using standard quality control measures and sample-specific maximum mitochondrial gene fraction, minimum number of genes per cell, minimum number of counts per cell, and maximum number of counts per cell were set to filter out low quality cells (Supplementary Table 1). To further filter out cells with low ATAC-seq quality, Muon was used to calculate ATAC-specific quality metrices. Sample-specific thresholds were identified for minimum and maximum number of counts, minimum and maximum TSS enrichment score as well as minimum and maximum nucleosome signal (Supplementary Table 1). *Doublet detection.* We used a combination of scrublet^72^ (v0.2.3), DoubletDetection^73^ (v4.2), scds^74^ (v1.10.0), scDblFinder^75^ (v1.11.4), DoubletFinder^76^ (v2.0.3) (default parameters, expected doublet rate 0.8) and SOLO^77^ (as implemented in scvi-tools^78^ v0.17.1) to detect doublets based on the gene expression modality. In addition, scDblFinder and its implementation of AMULET^79^ were used to identify doublets on the ATAC-seq modality. Cells consistently detected by three or more methods as doublets were excluded from further analysis.

#### Generation and quantification of common peak set

To merge the ATAC-seq data from individual samples, we followed the respective vignette on the Signac^80^ website. In brief, peaks from all samples were merged using the “reduce” function of the GenomicRanges (v1.46.1) package and only peaks on standard chromosomes were kept. Next, for each sample, fragment counts were determined using Signac and stored, together with gene expression data in a Seurat object, which were subsequently merged into a single object. *Normalization of ATAC-seq counts.* Signac was used to run TF-IDF normalization on ATAC-seq counts with default parameters. TF-IDF normalized count matrix was then imported into Muon. *Normalization of gene expression counts.* Prior to normalization, data from individual samples was merged. SCTransform^81^ (v0.3.3) was used for normalization using settings vst.flavor=”v2” and clip.range=c(-sqrt(n), sqrt(n)), where n represented the number of cells (n=27585).

#### Highly variable genes

The top 4,000 highly variable genes were identified using the devianceFeatureSelection function from the scry package^82^ (v1.6.0) with default parameters. *Label transfer.* To estimate initial cell type labels, we employed scANVI^83^ to transfer cell type annotations from an internal reference dataset. We first trained an scVI model (scvi-tools v 0.20.0), with parameters n_hidden=1024, n_latent=50, n_layers=2, gene_likelihood=’nb’, dispersion=’gene-batch’, sample names as batch key and the differentiation stage as categorial covariate, to generate a shared latent. Next, we trained a scANVI model and predicted cell type labels. Integration. To integrate the gene expression modality of the different Multiome samples, we trained new scVI model (scvi-tools v 0.20.0), with parameters n_hidden=1024, n_latent=50, n_layers=2, gene_likelihood=’nb’, dispersion=’gene-batch’ and sample names as batch key. We then trained a scANVI model using the predicted cell type labels. To integrate the chromatin accessibility modality, we used PoissonVI^84^, with parameters n_hidden=1024, n_latent=50, n_layers=2, sample names as batch key and the transferred labels as labels. To generate a joint neighborhood graph across modalities, we first computed k-nearest neighbor (KNN) graphs (k=16) on the latent spaces from both, scANVI and PoissonATAC, followed by the calculation of a weighted nearest neighborhood graph (WNN)^85^, combining the KNN graphs from both modalities. *Clustering and annotation.* Clustering was performed on the WNN graph using leiden^86^ clustering with resolution 2. To further separate subtypes of endocrine progenitors, the respective clusters were subclustered with lower resolutions (0.5-1). The resulting clusters were then annotated using a set of marker genes (Supplementary Table 1). For further analysis we focused on cells from the main endocrine and progenitor clusters, ignoring two minor clusters of unknown cell type.

### 7.4. Gene regulatory network inference

#### Peak to gene linking

To identify putative regulatory elements, we used the Signac (v1.9.0) LinkPeaks function with default parameters to calculate the correlation between chromatin accessibility and gene expression of nearby highly variable genes. *Motif detection.* Transcription factor motifs were detected in each peak using the AddMotifs function in Signac and the human_pwms_v2 motif collection provided by the chromVARmotifs package. *Feature selection.* We then selected the intersection between highly variable genes, expressed in more than 5% of cells, and transcription factors with matched motifs in the dataset as input genes for GRN inference. *GRN inference*. For every gene, we selected all linked peaks with a link score >= 0.25. We then calculated the spearman correlation between the peak accessibility and the expression of linked gene as well as the expression of any transcription factor with a motif within a given peak. We also calculated the spearman correlation between the linked gene and all transcription factors with motifs in the selected peaks. This results in a triplet of correlations for every possible regulatory interaction, which was used to further filter out meaningful target gene – peak – transcription factor relationships. An interaction was kept, if the absolute correlation between peak and transcription factor was >= 0.05 and the absolute correlation between target gene and transcription factor was >= 0.1.

### 7.5. Differential gene expression analysis

We used DElegate to calculate differentially expressed genes between each cell type and all other cell types to identify marker genes, as well as between Artemether treated and control. For the latter, we subset the data to the cell types of interest, filtered out genes detected in less than 5% of the cells, prior to differential expression analysis. We then applied induvial cut-offs for log fold change and adjusted p-value for every comparison, given by the minimum absolute log fold change where the corresponding z-score of the absolute log fold change was greater 0.5 and the minimum p-value where the corresponding z-score of the p-value was greater 0.25.

## 8. Proteomics

Cell pellets of the different treatments and replicates were resuspended in 25µl of 2x lysis buffer (Preomics GmbH, Martinsried) prior to tryptic digest applying the iST sample preparation kit (Preomics GmbH, Martinsried) following the manufacturer’s instructions. Dried peptides were resuspended in different volumes of 2% acetonitrile, 0.5% TFA, for normalization of varying cell counts in the cell pellets. Equal amounts of digested peptides were measured on a QExactive HF-X mass spectrometer (ThermoFisher Scientific) online coupled to a UItimate 3000 RSLC nano-HPLC (Dionex) as described in data-dependent acquisition mode (Grosche et al.,2016; Kaplan et al. 2022). Acquired raw files were analyzed in the Proteome Discoverer software (version 2.5, Thermo Fisher Scientific) for peptide and protein identification and quantification. Database search was performed using the Sequest HT search engine against the SwissProt Human database (Release 2020_02, 20432 sequences) as described (Kaplan et al. 2022). Protein abundances were calculated as the average of the 3 most abundant unique peptide group intensity values (TOP3) with a XCorr score >1. Protein abundance values were used for calculation of protein abundance ratios between groups and for ANOVA statistics (including Benjamini-Hochberg correction) within Proteome Discoverer 2.5. Peptide abundance values were normalized on total peptide amount. Match-between runs for label-free quantification was limited to a retention time window of 1 minute and a mass shift of 0.5 ppm. Missing values were imputed by low abundance resampling separately for each sample. Protein identifications and quantifications were exported and filtered for a false discovery rate <5%.

## 9. Statistics and reproducibility

No statistical method was used to pre-determine sample sizes, but the presented sample sizes are in line with previous publications. Statistical analysis and graphs were done using GraphPad Prism 9. Statistical methods, together with sample sizes and data representation information are provided in the respective figure legends. Data distribution was assumed to be normal, but not formally tested.

## Data availability

Source data will be provided upon publication. Any other data supporting the findings of this study are available from the corresponding author on reasonable request.

## Code availability

Jupyter notebooks to reproduce the analysis and figures will be made available on GitHub upon publication.

## Acknowledgements

We thank G. Lederer, G. Eckstein and S. Langer-Freitag for the karyotyping and the STR analysis. We thank the donor of the fibroblasts for supporting research projects with human material, Prof. Andreas Fritsche and his team for taking the skin samples. M. A. C is the recipient of a master and PhD fellowship granted by the Turkish Ministry of Education. This work was supported by the Helmholtz-Gemeinschaft (Helmholtz Portfolio Theme ‘Metabolic Dysfunction and Common Disease) and Deutsches Zentrum für Diabetesforschung (DZD). This project has received funding from the HumEN consortium funded by European Union’s Seventh Framework Program for Research, Technological Development and Demonstration under grant agreement no. 602889, the European Union’s Horizon 2020 research and innovation program under grant agreement ISLET number 874839.

