## Supplementary figures and tables for "Resolving human α *versus* β cell fate allocation for the generation of stem cell-derived islets"

### SUPPLEMENTAL INFORMATION

- Supplementary figures 1 to 5
- Supplementary table 1
- Material tables 1 to 6

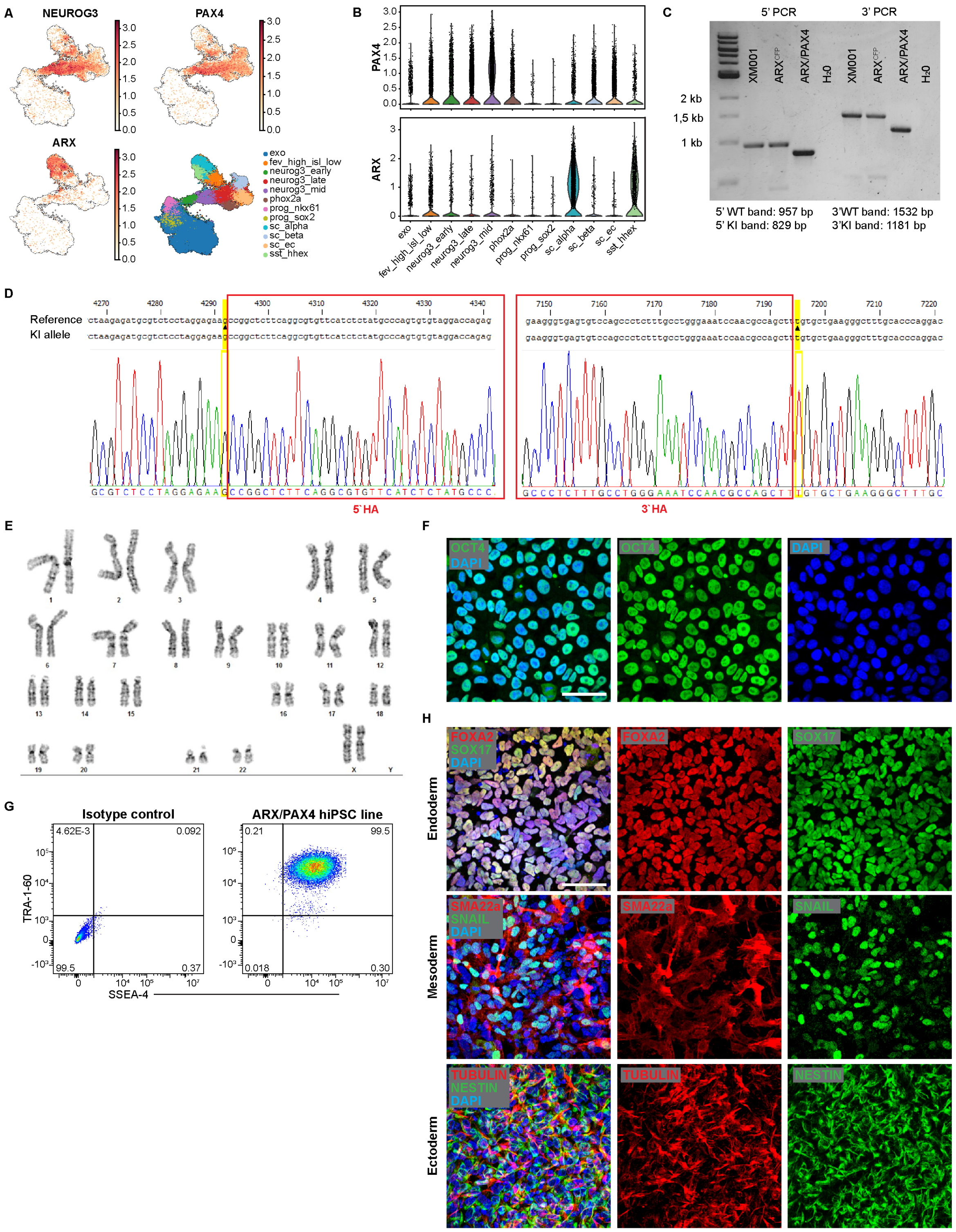


***Suppl. Fig. 1. Generation and quality controls of a human $ARX^{CFP/CFP}$/$PAX4^{mCherry/mCherry}$ reporter iPSC line.*** **(A-B)** UMAPs and violin plots of *NEUROG3, PAX4* and *ARX* expression in human SC-islets at differentiation stage 5 from the publicly available dataset from Veres *et al.*, 2019. **(C)** Genotyping of indicated iPSC clones for 5´ and 3´regions spanning the homology arms after transfection, sorting, and single-cell clonal culture. **(D)** Sanger sequencing of the 3' and 5' recombination boarders of the knock-in (KI) allele including the homology arms (HA). **(E)** Unperturbed female karyotype (46, XX) **(F-G)** Assessment of pluripotency using the nuclear markers OCT3/4 by IF (F), and the surface markers TRA-1-60 and SSEA-4 by flow cytometry analysis (G). **(H)** Direct differentiation toward the three germ layers (endoderm [FOXA2, SOX17], mesoderm [SM22-a, SNAIL] and ectoderm [TUBB3, NESTIN]) show full pluripotency and multilineage differentiation potential of selected clone. Scale bars 50 µm.

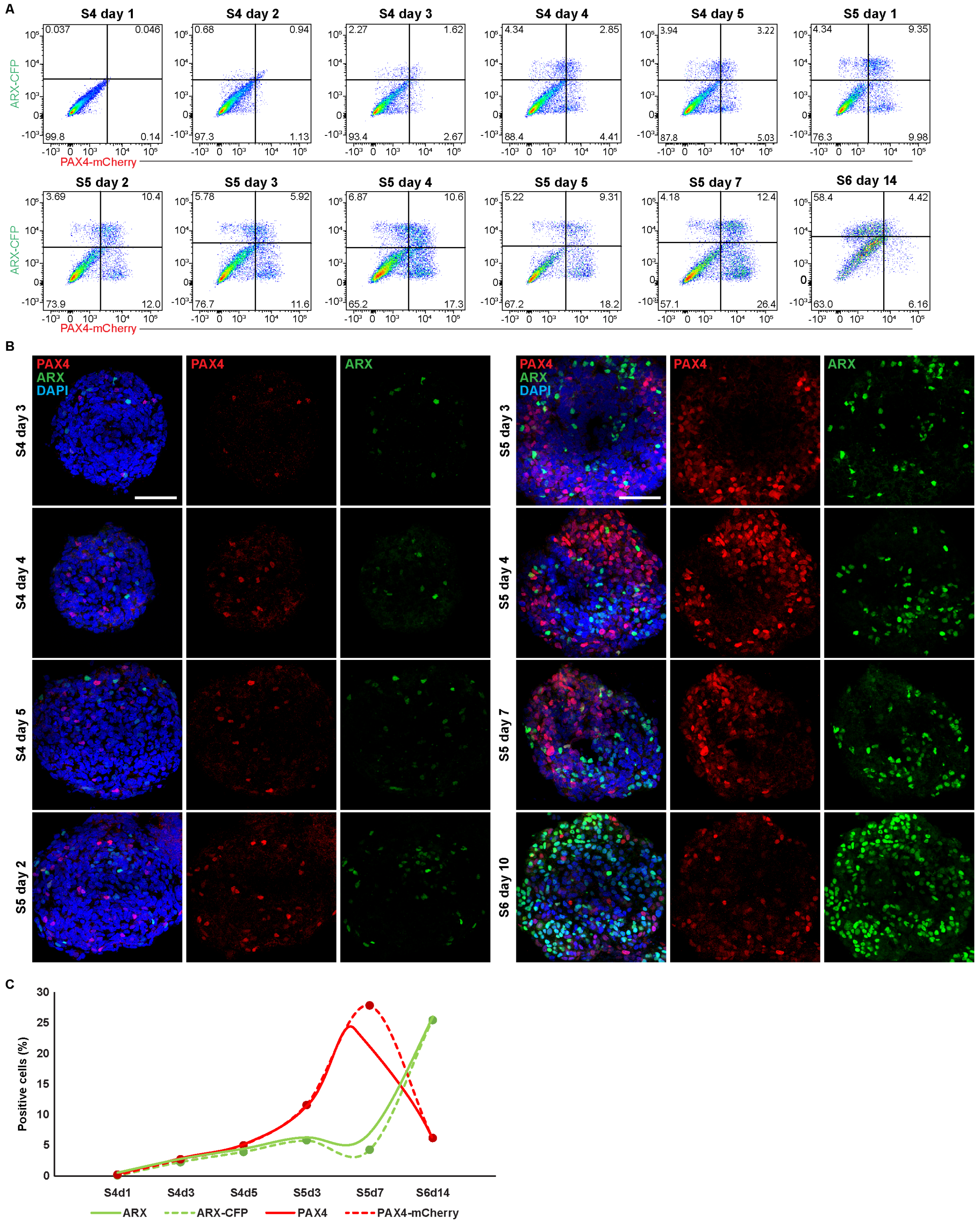


***Suppl. Fig. 2. Comparison between ARX-CFP, PAX4-mCherry, and endogenous proteins.*** **(A)** Flow cytometry analysis of the percentage of ARX-CFP and PAX4-mCherry positive cells starting from S4d1 (PP stage) to S6d14 (SC-β cells). **(B)** Representative maximum intensity projections of Z-stack confocal acquisitions of IF for endogenous ARX and PAX4 in wild-type clusters at the indicated stages. (C) Schematic representation of PAX4-mCherry (red dashed line), endogenous PAX4 (red solid line), ARX-CFP (green dashed line), and endogenous ARX (green solid line) dynamics during in vitro SC-β cells differentiation. Note the longer stability of PAX4-mCherry, making this reporter a short-term lineage tracer. n≥2 from distinct differentiation experiments.

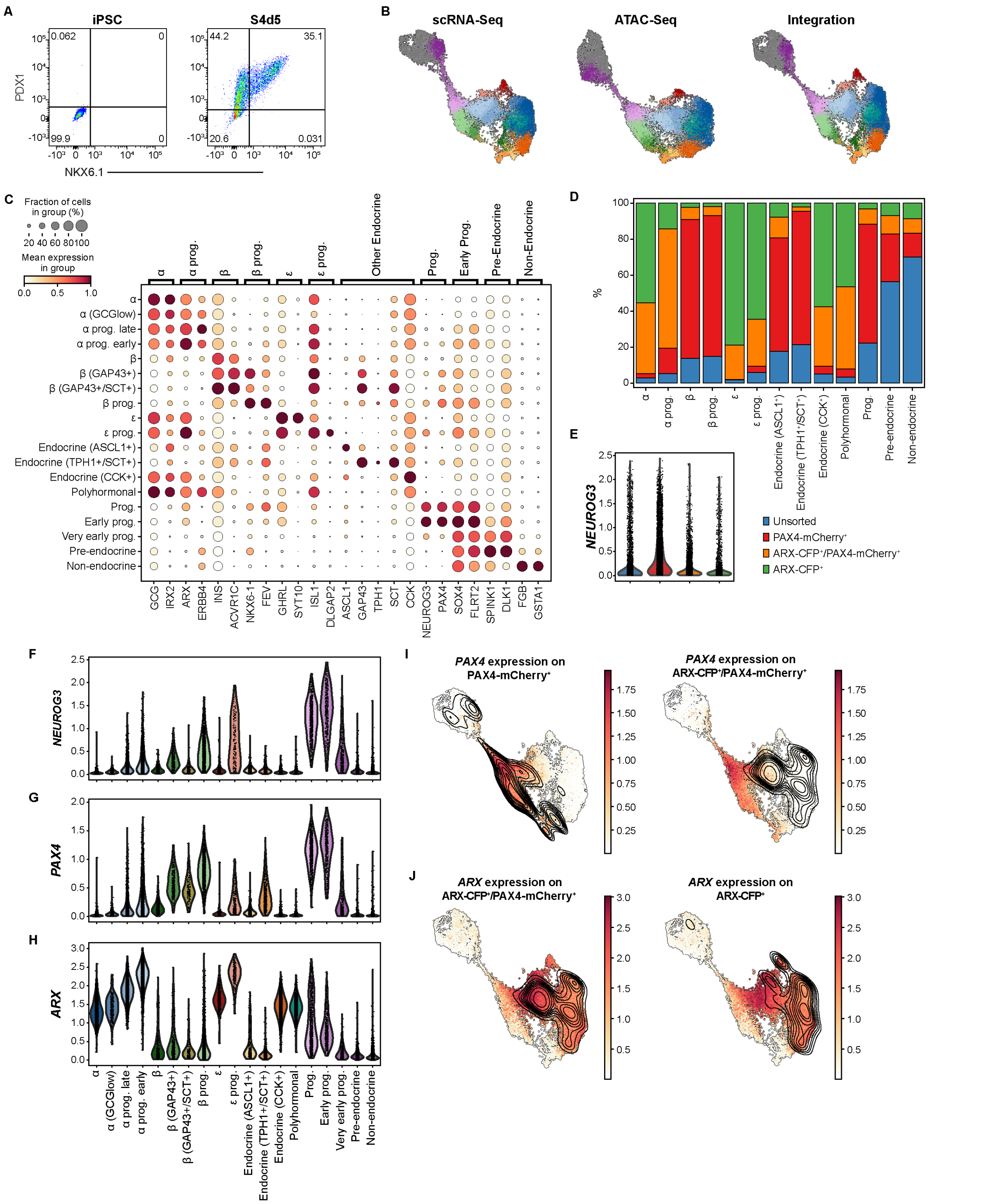


***Suppl. Fig. 3. Single cells multiomics suggests ARX repression of PAX4 during in vitro SC-β cells differentiation.*** **(A)** Flow cytometry plots showing differentiation efficiency (percentage of cells positive for PDX1 and NKX6.1) at S4d5 of samples subsequently submitted for scRNA-Seq and snATAC-Seq at S5d4 (S4d5) and iPSC controls. **(B)** UMAP embedding of single cells RNA-Sequencing (left), single cells ATAC-Sequencing (middle) and the integration of both (right) for all datasets. **(C)** Expression levels of annotated genes within all clusters. The color assigned to each gene in each cluster represents the average gene expression level, while the size of each circle represents the percentage of positive cells for each gene in that cluster. **(D)** Relative frequencies of cell types in each dataset as per annotated UMAP in fig. 2B. **(E)** Violin plots showing *NEUROG3* expression within all datasets. **(F-H)** Violin plots showing *NEUROG3, PAX4* and *ARX* expression within all cell clusters. **I-J)** *PAX4* and *ARX* expression UMAP including the embedding densities of PAX4-mCherry$^{+}$, ARX-CFP$^{+}$/PAX4-mCherry$^{+}$, and ARX-CFP$^{+}$ samples to illustrate temporal dynamics of TFs mRNA and their reporters.

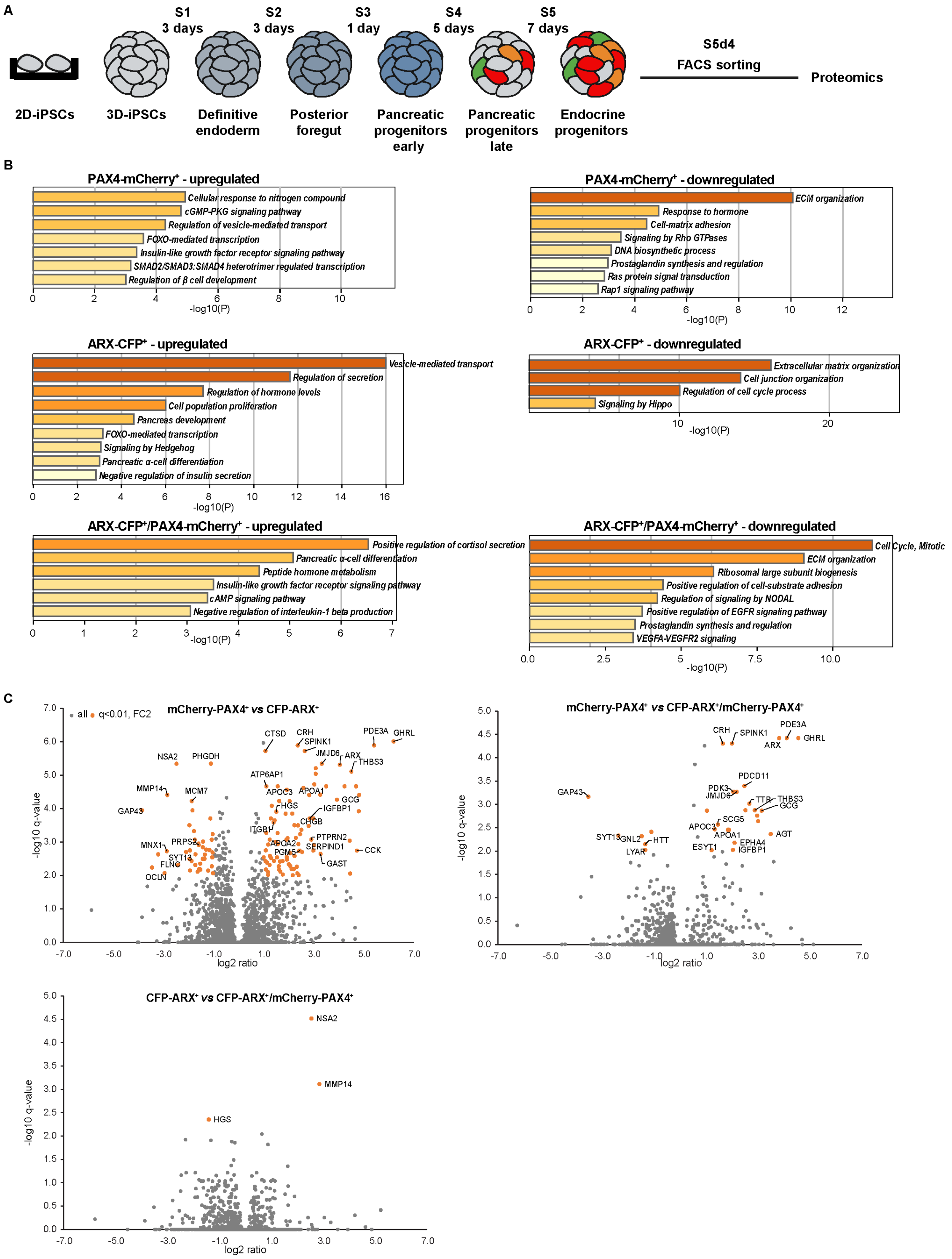


***Suppl. Fig. 4. Proteomics analysis confirms multiomic analysis.*** **(A)** Experimental strategy for proteomics analysis. **(B)** Ontology analysis of significant differentially expressed proteins for indicated samples against unsorted sample. **(C)** Volcano plots showing differentially expressed proteins between the specified samples. Note the small differences between the CFP-ARX⁺ and the CFP-ARX⁺/mCherry-PAX4⁺ samples. Proteins with a Foldchange of >2 fold (log2<-1 or log2>1) and a q-value >0.01 are represented by orange dots. q-values: Benjamini-Hochberg corrected p-values.

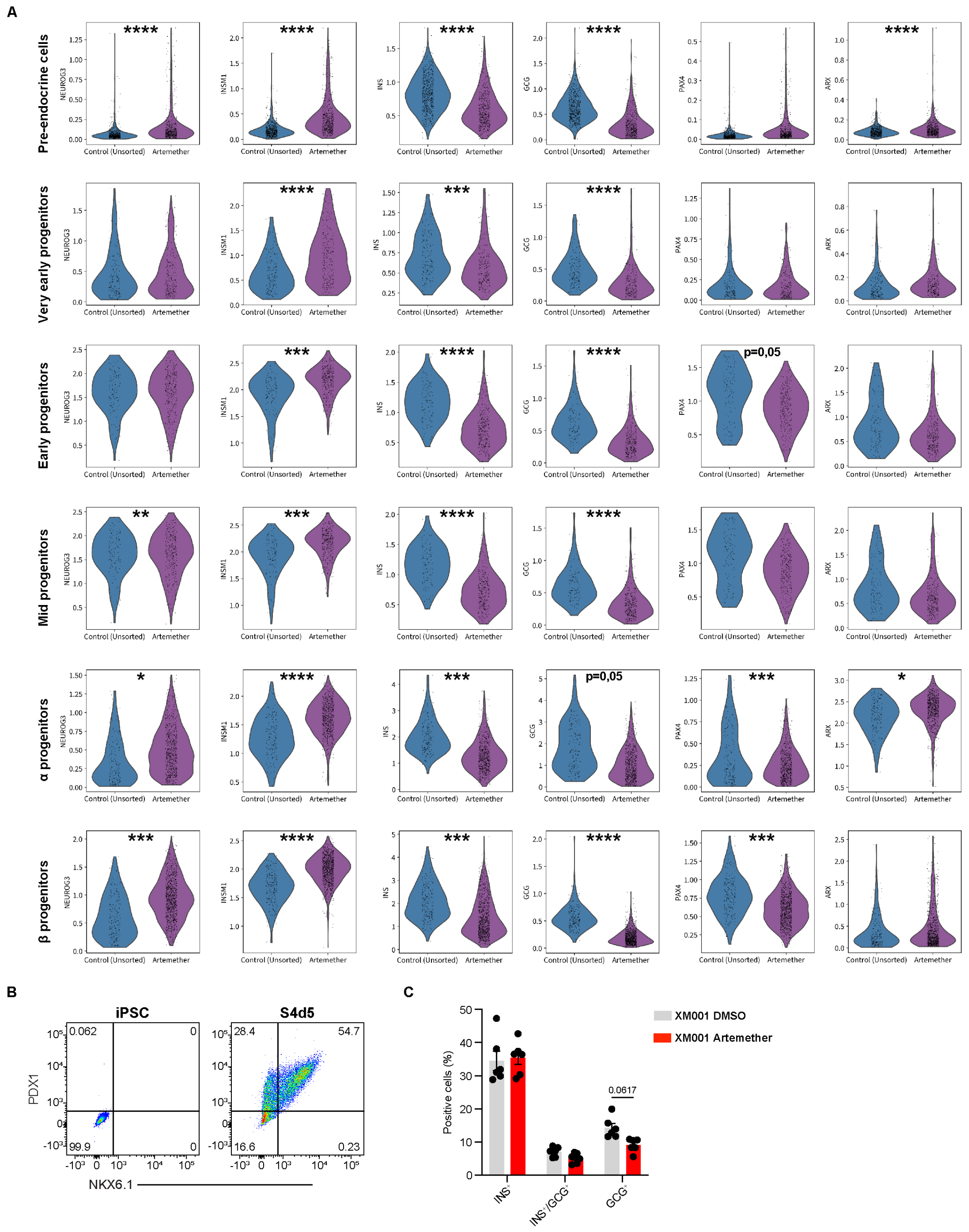


***Suppl. Fig. 5. Artemether promotes endocrine induction.*** **(A)** Violin plots showing *NEUROG3*, *INSM1*, *INS, GCG, PAX4*, and *ARX* expression within specified clusters of DMSO- and artemether-treated datasets at S5d4 **(B)** Flow cytometry plots showing differentiation efficiency (percentage of $PDX1^+$/$NKX6.1^+$ cells) at S4d5 of artemether-treated samples subsequently submitted for scRNA-Seq and snATAC-Seq at S5d4 (S4d5) and iPSC controls. (C) Quantification of hormone-positive cells at S6d14 after continuous treatment with artemether starting from S4. (G) after artemether treatment of wild-type hIPSC (XM001). Data are presented as mean ± SE. Two-ways ANOVA with Šídák multiple comparison test for C. * ≤ 0.05, ** ≤ 0.01, *** ≤ 0.001, **** ≤ 0.0001.

### Suppl. Table 1

| | GCG | ARX | INS | PAX4 | GAP43 | SCT | LMX1A | ASCL1 | TPH1 | CCK | GHRL | FEV | NEUROG3 | FLRT2 | SPINK1 | MKI67 | CHGA |
|---|---|---|---|---|---|---|---|---|---|---|---|---|---|---|---|---|---|
| Alpha | + | + | | | | | | | | | | | | | | | + |
| Alpha (GCG-low) | + | + | | | | | + | | | | | | | | | | + |
| Alpha Prog. early | | + | | | | | | | | | | | | | | | + |
| Alpha Prog. late | | + | | | | | | | | | | | | | | | + |
| Beta | | | + | | | | | | | | | | | | | | + |
| Beta (GAP43+) | | | + | + | + | | + | | | | | + | | | | | + |
| Beta (GAP43+/SCT+) | | | + | + | + | + | | | | | | | | | | | + |
| Beta Prog. | | | | + | | | + | | + | | | + | + | | | | + |
| Endocrine (ASCL1+) | | | | | | | | + | | | | + | | | | | + |
| Endocrine (TPH1+/SCT+) | | | | | + | + | | + | + | | | + | | | | | + |
| Endocrine. (CCK+) | + | + | | | | | | + | | + | | | | | | | + |
| Epsilon (GHRL+) | + | + | | | | | | | | | + | | | | | | + |
| Epsilon Prog. (GHRL+) | | + | | | | | | | | | + | | + | | | | + |
| Polyhormonal | + | + | | | | | | | | | | | | | | | + |
| Prog. | | | | + | | | + | | | | | + | + | + | | | + |
| Early Prog. | | | | + | | | + | | | | | | + | + | | | + |
| Early Prog. (Cycling) | | | | + | | | + | | | | | | + | + | | + | + |
| Very early Prog. | | | | | | | | + | | | | | + | + | + | + | |
| Progenitor primed | | | | | | | | | | | | | | + | + | + | |
| Non-endocrine | | | | | | | | | | | | | | | | | |
| Other Non-Endocrine (Removed) | | | | | | | | | | | | | | | | + | |
| Other Endocrine (Removed) | | + | | | | | + | | | | | | | | | | + |

**Material table 1. Primary antibody list**

| Name | Host/Conjugated | Company | Order number | Dilution IHC | Dilution FACS |
|---|---|---|---|---|---|
| **ARX** | Sheep | R&D Systems | AF7068 | 1:100 | |
| **Caspase-3, Cleaved** | Rabbit | Cell Signaling | 9661S | 1:100 | 1:100 |
| **FOXA2 (HNF-3β)** | Rabbit | Cell Signaling | 8186S | 1:300 | 1:200 |
| **GFP** | Chicken | Aves Labs | GFP-1020 | 1:300 | 1:300 |
| **Glucagon** | Mouse | Merck, Sigma-Aldrich | G2654-.2ML | 1:1000 | 1:500 |
| **Glucagon** | FITC | Novus Biologicals | NBP2-21803P | | 1:180 |
| **Insulin** | Guinea pig | Bio-Rad | 5330-0104G | 1:500 | 1:500 |
| **Insulin-T56-706** | APC | BD | 565689 | | 1:40 |
| **Ki-67** | Rabbit | Abcam | ab15580 | 1:300 | 1:500 |
| **RFP (5F8)** | Rat | chromotek | ORD003515 | 1:300 | 1:300 |
| **NEUROG3** | Rabbit | Aviva Systems Biology | OACD05949 | 1:200 | 1:100 |
| **NESTIN (10C2)** | Mouse | Abcam | ab22035 | 1:200 | |
| **NKX2-2** | Mouse | Abcam | ab187375-500ul | 1:300 | 1:200 |
| **NKX6-1** | Rabbit | Bio-techne, Novus | NBP1-82553 | 1:300 | 1:200 |
| **Oct-3/4** | Goat | Santa Cruz | sc-8628 | 1:500 | |
| **PDX1** | Goat | R&D Systems | AF2419 | 1:300 | 1:100 |
| **PAX4** | Rabbit | Life Technologies | PA1-108 | 1:200 | |
| **SOX17** | Goat | Neuromics | GT15094 | 1:400 | 1:200 |
| **SLC18A** | | Atlas Antibodies | ATAHPA063797-100 | | |
| **SM22 alpha (Transgelin)** | Rabbit | Abcam | ab14106 | 1:100 | |
| **Somatostatin (H-11)** | Mouse | Santa Cruz | sc-74556 | 1:300 | |
| **Snail** | Goat | R&D Systems | AF3639 | 1:300 | |
| **Tubulin beta III** | Rabbit | Abcam | ab18207 | 1:1000 | |

**Material table 2. Secondary antibody list**

| Name | Host | Fluorophore | Company | Order number | Dilution IHC/FACS |
|---|---|---|---|---|---|
| **anti-Guinea Pig IgG** | Donkey | AlexaFluor 488 | Biozol, Jackson ImmunoResearch | 706-545-148 | 1:500 |
| **anti-Guinea Pig IgG** | Donkey | AlexaFluor 647 | Biozol, Jackson ImmunoResearch | 706-165-148 | 1:500 |
| **anti-Goat IgG** | Donkey | AlexaFluor 488 | Thermo Fisher Scientific, Invitrogen | A11055 | 1:500 |
| **anti-Goat IgG** | Donkey | AlexaFluor 555 | Thermo Fisher Scientific, Invitrogen | A21432 | 1:500 |
| **anti-Goat** | Donkey | AlexaFluor 647 | Biozol, Jackson ImmunoResearch | 705-605-147 | 1:500 |
| **anti-Mouse IgG** | Donkey | AlexaFluor 488 | Thermo Fisher Scientific, Invitrogen | A21202 | 1:500 |
| **anti-Mouse IgG** | Donkey | AlexaFluor 555 | Thermo Fisher Scientific, Invitrogen | A31570 | 1:500 |
| **anti-Mouse** | Donkey | AlexaFluor 647 | Thermo Fisher Scientific, Invitrogen | 715-605-151 | 1:500 |
| **anti-Rabbit IgG** | Donkey | AlexaFluor 488 | Thermo Fisher Scientific, Invitrogen | A21206 | 1:500 |
| **anti-Rabbit IgG** | Donkey | AlexaFluor 555 | Thermo Fisher Scientific, Invitrogen | A31572 | 1:500 |
| **anti-Rabbit IgG** | Donkey | AlexaFluor 647 | Thermo Fisher Scientific, Invitrogen | A31573 | 1:500 |
| **anti-Rat IgG** | Donkey | Cy3 | Biozol, Jackson ImmunoResearch | 712-165-153 | 1:500 |
| **anti-Sheep IgG** | Donkey | AlexaFluor 488 | Biozol, Jackson ImmunoResearch | 713-546-147 | 1:500 |

**Material table 3. PAX4 single guide RNA**

| PAX4-sgRNA | Sequence |
|---|---|
| **Forward PAX4-sgRNA** | CACC GGG GAGCATGCATCAGGACGGTG |
| **Reverse PAX4-sgRNA** | AAACCACCGTCCTGATGCATGCTC CCC |
| **Kozak sequence** | CACC |
| **PAM sequence** | 5'-NGG-3' |

**Material table 4. Primer for genotyping**

| Primer | Sequence | Primer number |
|---|---|---|
| **Pax4 3' KI forward primer** | GGATCACTCTCGGCATGGAC | EP308 |
| **Pax4 3'KI reverse primer** | TCTGAGGGCTTCTGGGACTTGG | EP1849 |
| **Pax4 5'KI forward primer** | CAGCAGGTTAGAGATGCTAAGAG | EP408 |
| **Pax4 5'KI reverse primer** | GCCTTCTCAGCCCTGGAAGACAC | EP1848 |

**Material table 5. Primer for cloning homology arms of Pax4**

| Primer | Sequence with overhangs | Tm |
|---|---|---|
| **5'HA Pax4 forward** | CTATAGGGCGAATTGGAGCTCCACCGC GCCGGCTCTTCAGGCGTGTTCATCTC | 73°C |
| **5'HA Pax4 reverse** | CTTCGCTGGCTCTGGCATGGTGGCGC GGCTGACCCTCCTCAGAAGGATGAGAC | 71°C |
| **3'HA Pax4 forward** | GCGACGTTGAGGAAAACCCAGGACCAATGCATCAGGACGGTGAGGAGCCTGGG | 74°C |
| **3'HA Pax4 forward** | CCCCTCGAGGTCGACGGTATCGATA GCTGGCGTTGGATTTCCCAGGCAAAGAGG | 75°C |

| Material table 6. Kit | Company | Catalogue number |
|---|---|---|
| **Gel Extraction Kit** | Qiagen | 28704 |
| **PCR Purification Kit** | Qiagen | 28104 |
| **Plasmid Mini Kit** | Qiagen | 12125 |
| **Plasmid PLUS Midi Kit** | Qiagen | 12943 |
| **Insulin ELISA** | Mercodia | 10-1113-01 |
